# Hybridization chain reaction variants with enhanced sensitivity for detecting challenging mRNA targets

**DOI:** 10.1101/2025.09.09.675218

**Authors:** Chanpreet Singh, Namrata Bali, Gerard M. Coughlin, Jin Xu, Juni Y Polansky, Ulrich Herget, Madelyn S. Gilbert, Tasha Cammidge, Giada Spigolon, Yelena Smirnova, Viviana Gradinaru, Kai Zinn, David A. Prober

## Abstract

Compared to traditional enzyme-based *in situ* amplification methods, Hybridization Chain Reaction v3.0 (HCR v3.0) offers high specificity for spatial RNA visualization but lacks the sensitivity required to robustly detect short or low-abundance targets, particularly in thick tissue with high autofluorescence. Here, we describe three HCR variants that combine the specificity of HCR v3.0 with additional signal amplification through catalytic reporter deposition (HCR-Cat), immunostaining (HCR-Immuno), or iterative HCR and immunodetection (HCR-Multi). These methods substantially enhance detection sensitivity, enabling robust spatial visualization of low-abundance transcripts, improved performance in challenging tissue environments, and compatibility with fluorescence- and alkaline phosphatase-based chromogenic detection. These methods enable transcript-resolution imaging, even when using a limited number of probes, thereby expanding the range of biological applications accessible to HCR.

**Summary Statement:** Enhanced-sensitivity HCR variants enable robust spatial detection of low-abundance and difficult-to-detect RNA targets while preserving the specificity of standard HCR.

## Introduction

Spatial biology has revolutionized our understanding of cellular processes by providing crucial insights into the tissue-specific organization and regulation of gene expression. In particular, spatial RNA profiling enables precise mapping of RNA expression patterns within their native cellular environment. Hybridization Chain Reaction v3.0 (HCR v3.0) (Choi et al., 2018) enables multiplexed, quantitative, high-resolution imaging of RNA targets with low background through the use of split-initiator probes. The amplified signal scales approximately linearly with both the number of probes targeting each RNA and the number of RNA target molecules. Although increasing the number of probes can improve sensitivity, this is not feasible for short transcripts or targets that have only a limited number of specific probe-binding sites. Consequently, detection sensitivity can be limited for short transcripts, low-abundance targets, or experiments using sparse probe sets. These challenges are further exacerbated in thick tissue samples with high levels of autofluorescence. Here, we describe three HCR variants — HCR-Cat, HCR-Immuno, and HCR-Multi — that enhance detection sensitivity while preserving the specificity and multiplexing capabilities of HCR v3.0. These methods expand the range of targets and applications that are accessible to HCR, allowing robust detection of RNA targets that were previously challenging to detect.

## Results

### HCR-Cat enhances mRNA detection sensitivity in whole-mount zebrafish larvae

Hybridization Chain Reaction (HCR) v3.0 provides highly specific spatial RNA detection through split-initiator probes but can fail to detect transcripts that are short, expressed at low levels, or present in highly autofluorescent tissues. To overcome these limitations, we developed HCR-Cat (Hybridization Chain Reaction with Catalytic reporter deposition), in which conventional fluorophore-conjugated HCR amplifiers are replaced with hapten-conjugated amplifiers that are recognized by enzyme-conjugated antibodies (**Fig. 1A,B**). Following HCR amplification, horseradish peroxidase (HRP)-conjugated antibodies catalyze deposition of fluorescent tyramide reporters, substantially increasing signal intensity while retaining the target specificity provided by split-initiator HCR probes. To evaluate HCR-Cat, we first examined expression of the neuropeptide hypocretin (*hcrt*) (Prober et al., 2006) in whole-mount larval zebrafish at 5 days post fertilization (5 dpf). HCR-Cat, performed using fluorescein isothiocyanate (FITC)-conjugated HCR amplifiers and an HRP-conjugated anti-Fluorescein (anti-Flu) antibody, generated substantially stronger fluorescence than HCR v3.0 across the full range of laser powers tested while maintaining the same detector gain (**Fig. 1C–E”**, **Fig. S1A–B”**). While HCR v3.0 produced relatively low levels of fluorescence that increased gradually with increasing laser power, HCR-Cat generated much brighter fluorescence that was readily detectable even at low laser powers. Quantification demonstrated that HCR-Cat produced a more than two-orders-of-magnitude increase in net fluorescence signal (i.e., total fluorescence - background fluorescence) compared with HCR v3.0 (∼240-fold increase averaged across all laser powers tested) (**Fig. 1F**), with signal approaching saturation at approximately 50% laser power. Background fluorescence also increased modestly at higher laser powers (**Fig. 1F’**), but the increase in target signal was substantially greater, resulting in markedly improved net signal.

**Fig. 1:**
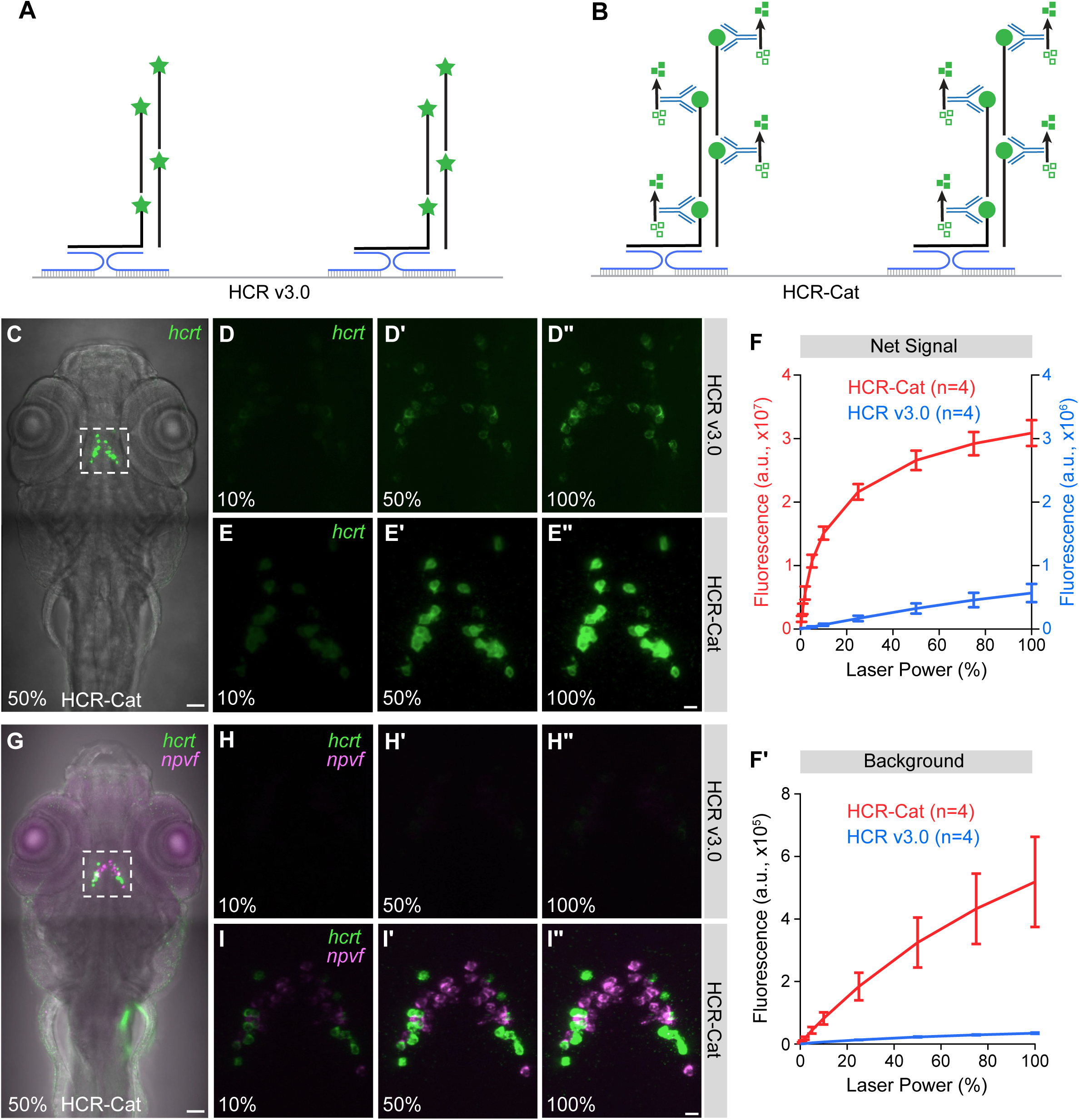
HCR-Cat enhances mRNA detection sensitivity compared to HCR v3.0 in whole-mount zebrafish larvae. **(A–B)** Split probe pairs (blue) hybridize to their target RNA (gray), forming a complete initiator that triggers the chain-reaction assembly of HCR amplifier hairpins (black). Compared to Alexa Fluor-conjugated HCR v3.0 amplifiers (green stars) **(A)**, HCR-Cat amplifiers **(B)** are conjugated to haptens (green circles) that are detected by an HRP-conjugated antibody (blue Y), which catalytically deposits TSA reporter molecules (open green squares converted to closed green squares). **(C)** Detection of *hcrt* with 8 probes using HCR-Cat at 50% laser power. **(D–E”)** Detection of *hcrt* with 8 probes using HCR v3.0 (**D–D”**) and HCR-Cat (**E– E”**) at different laser powers. **(F)** Quantification of net signal intensity (i.e., total fluorescence - background fluorescence) using 8 *hcrt* probes and either HCR v3.0 (blue, n=4 fish) or HCR-Cat (red, n=4 fish). HCR-Cat showed significantly higher signal compared to HCR v3.0 at all laser powers (p < 0.05 at 1% laser power, p < 0.0001 for other laser powers). There was no significant difference in HCR-Cat signal between 50% and higher laser powers, indicating that the net signal approached saturation at 50% laser power. **(F’)** Quantification of background fluorescence used to calculate net signal in **(F)**. HCR-Cat showed significantly higher background than HCR v3.0 beginning at 25% laser (p < 0.05), with highly significant differences at all higher laser powers (p < 0.0001). For each sample, the boxed region in **(C)** was imaged by gradually increasing the laser power from 0% to 100% while maintaining a fixed detector gain. Mean ± SEM (arbitrary units, a.u.) are shown (**F, F’**). **(G)** Co-detection of *hcrt* and *npvf* with 8 and 10 probes, respectively, using HCR-Cat at 50% laser power for both channels. **(H–I”)** Co-detection of *hcrt* and *npvf* with 8 and 10 probes, respectively, showed higher signal for both targets using HCR-Cat (**I–I”**) compared to HCR v3.0 (**H–H”**). A higher digital gain was used to detect *hcrt* in (**C–F”**) than in (**G–I”**). All experiments used at least 4 fish. High-magnification images shown in (**D–E”**) and (**H–I”**) correspond to the boxed regions in (**C,G**) and **Fig. S1** (**A”– D”**). Representative images shown are maximum intensity projections of z-stacks across the entire cell populations. Alexa Fluor 488 (**D, H**), Alexa Fluor 546 (**H**), TSA-Flu (**E, I**), and TSA-Cy3 (**I**) were used. Scale bars: 50 μm (**C, G**) and 10 μm (**E”, I”**). Statistical significance was assessed using Two-way ANOVA with amplification method and laser power as factors, followed by Tukey’s multiple-comparisons test.

To determine whether the improved sensitivity of HCR-Cat generalized beyond *hcrt*, we evaluated several additional neuronal transcripts in whole-mount zebrafish larvae. HCR-Cat similarly enhanced detection of *dopamine* β*-hydroxylase* (*dbh*) (Prober et al., 2006), *dopamine transporter* (*dat*) (Chen et al., 2009), and *vasoactive intestinal peptide* (*vip*) (Chen et al., 2017), compared to HCR v3.0 (**Figs. S2A–E’’** and **S3**), demonstrating that the improved sensitivity is broadly applicable across multiple transcript targets. Increasing the digital gain used to image HCR v3.0 samples increased signal visibility but did not reproduce the fluorescent levels observed with HCR-Cat under the conditions tested (**Fig. S2C–C’’**, **E–E’’**). A comparable increase in net signal was obtained using Digoxigenin (DIG)-conjugated HCR amplifiers and an HRP-conjugated anti-DIG antibody to detect *neuropeptide VF* (*npvf*) (Lee et al., 2017), which is expressed in hypothalamic neurons that are co-mingled with *hcrt*-expressing neurons (**Fig. S4**). Net signal increased by approximately 280-fold relative to HCR v3.0 (averaged across all laser powers tested) (**Fig. S4E,E’**), similar to the ∼240-fold increase that we observed for *hcrt*, demonstrating that HCR-Cat performs similarly with FITC- and DIG-based detection chemistries. Finally, simultaneous detection of *hcrt* and *npvf* using FITC- and DIG-conjugated amplifiers, respectively, enabled robust multiplex transcript detection within the same specimen (**Fig. 1G–I’’**, **Fig. S1C–D’’**), demonstrating that independent hapten-antibody detection systems can be combined for multiplex transcript detection using HCR-Cat.

### HCR-Cat improves transcript detection across tissues, tissue volumes, and species

To determine whether the improved sensitivity of HCR-Cat extends beyond whole-mount zebrafish larvae, we next evaluated its performance in 100 μm-thick mouse brain tissue slices. HCR-Cat robustly detected the regionally restricted neuronal markers dopamine receptor D1 (*Drd1*) (Ogata et al., 2022), dopamine receptor D2 (*Drd2*) (Ogata et al., 2022), tachykinin receptor 1 (*Tacr1*) (Allen Institute for Brain Science, 2004), tachykinin receptor 3 (*Tacr3*) (Allen Institute for Brain Science, 2004), and tyrosine hydroxylase (*Th*) (Gupta et al., 1990) at laser powers that failed to produce detectable signal using HCR v3.0 (**Fig. 2A**). The expression patterns observed for these targets were consistent with previously reported spatial expression patterns and with chromogenic *in situ* hybridization data from the Allen Brain Atlas (Allen Institute for Brain Science, 2004). Similar enhancement was observed for the pan-neuronal markers synaptosomal-associated protein 25 (*Snap25*) (Allen Institute for Brain Science, 2004) and enolase 2 (*Eno2*) (Allen Institute for Brain Science, 2004), as well as the glial marker solute carrier family 1 member 3 (*Slc1a3*) (Allen Institute for Brain Science, 2004) (**Fig. S5A**), demonstrating that the improved sensitivity is broadly applicable across multiple neuronal and glial transcript targets.

**Fig. 2:**
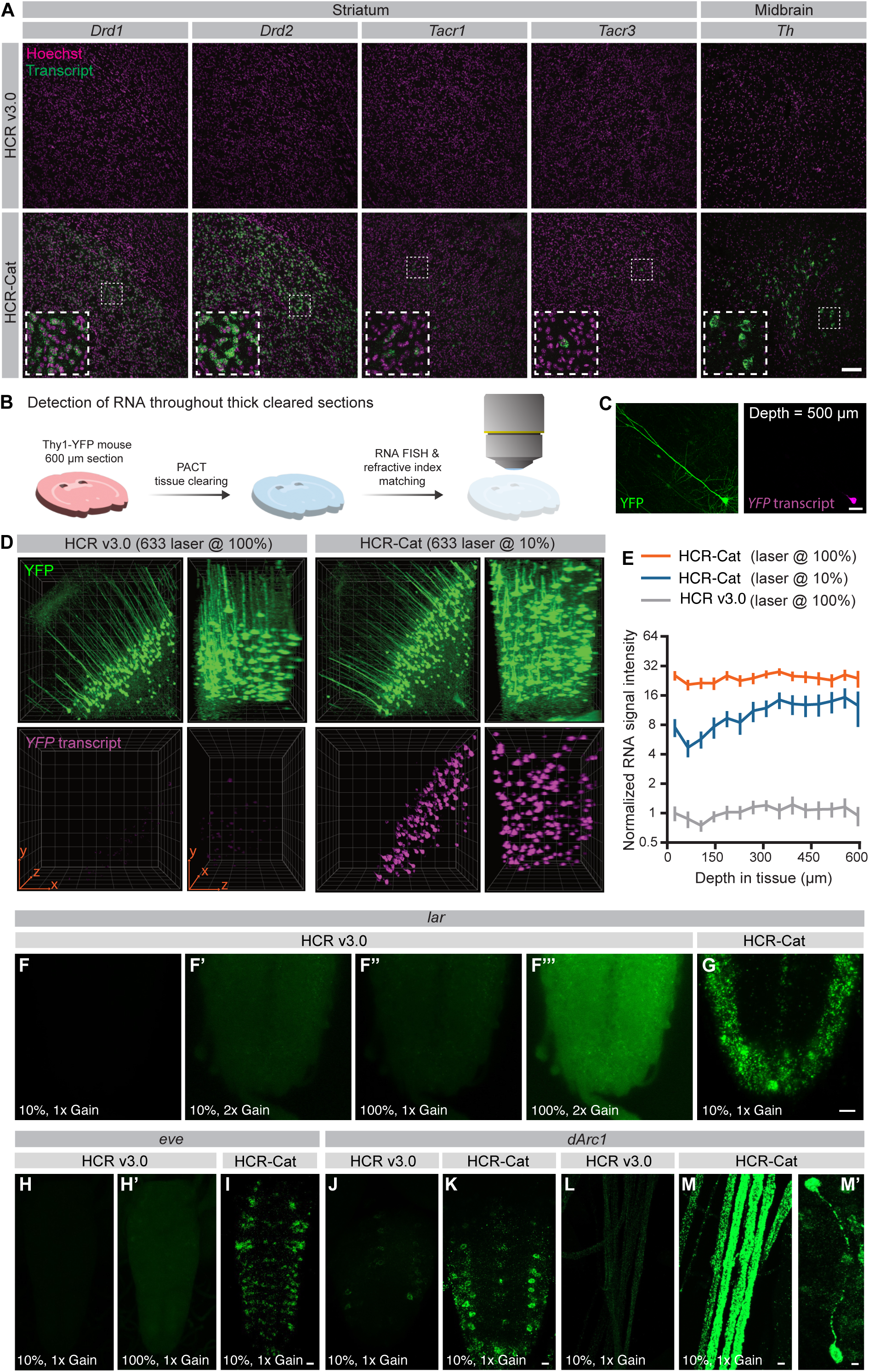
HCR-Cat enhances mRNA detection in mouse brain and whole-mount *Drosophila* larvae. **(A)** Comparison of HCR-Cat with HCR v3.0 for detection of neuron subtype markers in the mouse striatum and midbrain. Consistent with known expression patterns, HCR-Cat detected *Drd1* (15 probes)*-* and *Drd2* (16 probes)*-*positive neurons throughout the striatum, whereas *Tacr1* (13 probes) and *Tacr3* (13 probes) labeled sparser populations of cells. Cell-specific expression of *Th* (13 probes) was detected in the ventral tegmental area and substantia nigra of the midbrain. Each staining was performed on 100 μm-thick sections from 3 mice. Representative images shown are maximum intensity projections of z-stacks. Boxed regions are shown at higher magnification in each inset. **(B)** Experimental paradigm for detection of mRNA throughout thick sections of Thy1-YFP mouse brain using PACT clearing, RNA labeling with HCR v3.0 or HCR-Cat, followed by refractive index matching and imaging. **(C)** Representative YFP-positive neuron at 500 μm depth, showing strong and specific *YFP* (7 probes) mRNA signal using HCR-Cat. **(D)** Confocal stacks through 600 μm of PACT-cleared Thy1-YFP mouse cortex, showing detection of *YFP* transcript using HCR-Cat but not HCR v3.0. HCR v3.0 samples were imaged at 100% laser power, whereas HCR-Cat samples were imaged with 10% laser power. No correction for imaging depth was applied. Representative images from 4 mice are shown. **(E)** Quantification of *YFP* mRNA signal intensity in cell bodies segmented using the YFP channel. Data were normalized to the HCR v3.0 mRNA signal intensity at the tissue surface. HCR-Cat yielded an approximately 25-fold increase in *YFP* mRNA signal compared to HCR v3.0 when imaged at the same laser power (p < 0.0001, at all tissue depths). Decreasing the laser power to 10% still yielded an approximately 10-fold increase in *YFP* mRNA signal compared to HCR v3.0 (p < 0.05 at all depths except 64.6 μm and 105.4 μm). Mean ± 95% confidence interval from 17-63 cells per depth, pooled from 4 mice, is shown. Statistical significance was tested with a mixed effects analysis with Dunnett’s multiple comparison test, treating the animal as the unit of replication. Complete statistical results, including adjusted p-values for pairwise comparisons, are provided in Table S1. **(F–G)** Comparison of HCR-Cat and HCR v3.0 for detection of *lar* in the larval *Drosophila* ventral nerve cord (VNC). Using 20 probes, HCR-Cat produced a robust signal at 10% laser power, whereas HCR v3.0 failed to yield a clear signal even at 100% laser power and double the detector gain. **(H–I)** Comparison of HCR-Cat and HCR v3.0 for detection of *eve* in the larval *Drosophila* VNC. Using 19 probes, HCR-Cat produced a robust signal at 10% laser power, whereas HCR v3.0 failed to yield detectable signal even at 100% laser power. **(J–K)** Comparison of HCR-Cat and HCR v3.0 for detection of *dArc1* in the larval *Drosophila* VNC. HCR-Cat dramatically increased the signal compared to HCR v3.0 when imaged using the same laser power and detector gain. **(L–M)** Comparison of HCR-Cat and HCR v3.0 for detection of *dArc1* in motor nerves (**M**, bundles of multiple projections are shown) and individual neuronal soma and projections (**M’**) of the larval *Drosophila* brain. Using 19 probes, HCR-Cat produced a robust signal compared to HCR v3.0. Each experiment used at least 7 *Drosophila* larvae. Representative images shown are maximum intensity projections of z-stacks across the entire cell populations. Alexa Fluor 488 (**A**, top panel**; F–F’”, H–H’, J, L**), Alexa Fluor 647 (**D**, left two panels), TSA-Flu (**G, I, K, M–M’**), TSA-Alexa Fluor 488 (**A**, bottom panel) and TSA-Alexa Fluor 647 (**D**, right two panels) were used. Scale bars: 100 μm (**A**), 50 μm (**C**), 160 μm (**D**, orange arrows in plane of image) and 10 μm (**F–M’**).

We next asked whether HCR-Cat could improve transcript detection throughout large tissue volumes. Using 600 μm-thick PACT-cleared (Treweek et al., 2015; Yang et al., 2014) brain sections from Thy1-YFP mice (Feng et al., 2000), we detected *YFP* transcript following tissue clearing, refractive index matching, and volumetric imaging (**Fig. 2B**). The sparse expression of YFP protein enabled segmentation of individual pyramidal neurons for quantitative analysis. HCR-Cat generated robust and specific *YFP* transcript labeling throughout the cleared tissue volume, whereas HCR v3.0 produced only weak fluorescence despite imaging at ten-fold higher laser power (**Fig. 2C,D**). Quantification revealed an approximately 25-fold increase in net signal with HCR-Cat compared to HCR v3.0 that was maintained throughout the tissue without signal attenuation at deeper sections (**Fig. 2E**). Comparable enhancement was observed for the endogenous neuronal marker *Snap25*, which exhibited an approximately 12-fold increase in signal relative to HCR v3.0 (**Fig. S5B,C**).

To determine whether HCR-Cat is similarly effective in other large vertebrate tissues, we next examined whole adult zebrafish brain following iDISCO+ tissue clearing (Kenney et al., 2021). HCR-Cat generated bright and spatially restricted labeling of *dbh*-expressing neurons, whereas HCR v3.0 failed to produce detectable signal under the same imaging conditions (**Fig. S2F,G**), demonstrating that the enhanced sensitivity of HCR-Cat is maintained in intact adult vertebrate brain.

We also evaluated HCR-Cat in larval *Drosophila* tissues. HCR-Cat dramatically enhanced detection of the low-abundance transcripts *leukocyte antigen-related-like* (*lar*) (Bali et al., 2022) and *even-skipped* (*eve*) (Srinivasan et al., 2012) compared to HCR v3.0 (**Fig. 2F–I**). Whereas HCR v3.0 failed to detect *eve* even at maximum laser power and produced only weak signal for *lar*, HCR-Cat generated robust labeling under identical imaging conditions. We further applied HCR-Cat to *Drosophila activity-regulated cytoskeleton-associated protein 1* (*dArc1*) (Ashley et al., 2018) and observed strong transcript signal in ventral nerve cord motor neurons, consistent with previous genetic studies that inferred motor neuron expression (Ashley et al., 2018) (**Fig. 2J–M**). HCR-Cat also detected *dArc1* mRNA within motor nerve projections and in the axonal processes of a small population of neurons in the brain (**Fig. 2M,M’**), providing the first direct visualization of *dArc1* transcript localization within these neuronal processes.

### HCR-Immuno and HCR-Multi enhance mRNA detection sensitivity while preserving spatial resolution

Although HCR-Cat dramatically increases detection sensitivity, catalytic deposition of fluorescent tyramide reporters results in signal diffusion that limits subcellular resolution. To avoid this limitation while maintaining enhanced sensitivity, we developed HCR-Immuno (Hybridization Chain reaction with Immunofluorescence detection), in which hapten-conjugated HCR amplifiers are detected by primary antibodies followed by fluorophore-conjugated secondary antibodies (**Fig. 3A,B**). Unlike catalytic reporter deposition, antibody labeling remains localized to the HCR amplifiers, thereby preserving the spatial resolution of the HCR signal. Using *npvf* as a test transcript in larval zebrafish, HCR-Immuno generated substantially stronger fluorescence than HCR v3.0 across the range of laser powers tested (**Fig. 3C–D’’**). Quantification revealed an approximately 4.5-fold increase in net signal compared to HCR v3.0 (averaged across all laser powers tested) (**Fig. 3E**). Although background fluorescence increased modestly at higher laser powers (**Fig. 3E’**), the increase in target signal was substantially greater, resulting in significantly improved net signal.

**Fig. 3:**
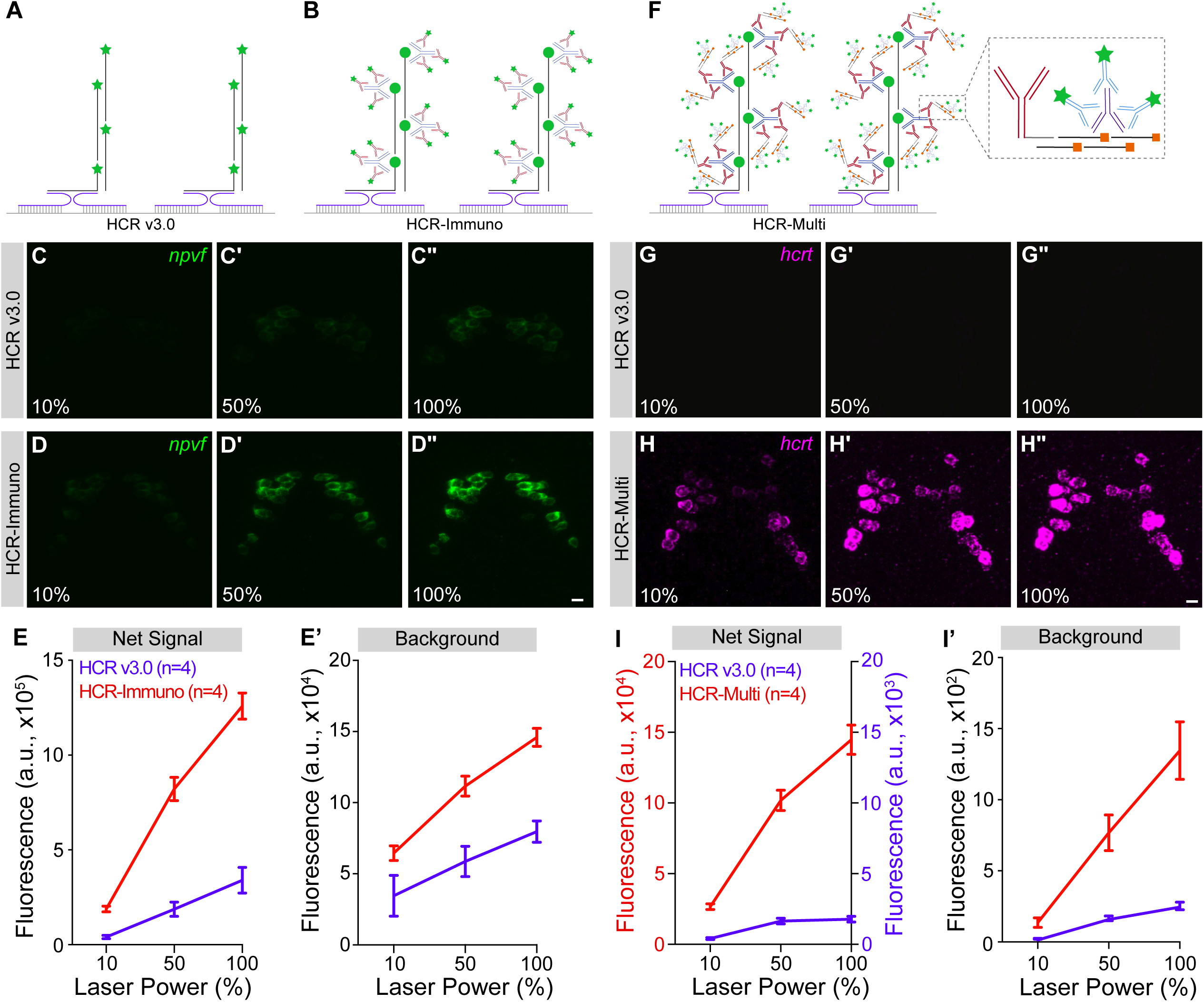
HCR-Immuno and HCR-Multi enhance mRNA detection sensitivity in whole-mount zebrafish larvae compared to HCR v3.0. **(A–B)** Split probe pairs (blue) hybridize to their target RNA (gray), forming a complete initiator that triggers the chain-reaction assembly of HCR amplifier hairpins (black). Compared to Alexa Fluor-conjugated (green stars) HCR v3.0 amplifiers (**A**), HCR-Immuno amplifiers (**B**) are conjugated to haptens (green circles) that are detected by a primary antibody (blue Y), followed by detection using a secondary antibody (red Y) conjugated to Alexa Fluor fluorophores (green stars). HCR-Immuno produced a robust signal for *npvf* (**D–D”**) compared to HCR v3.0 (**C–C”**) using 10 probes. **(E)** Quantification of net signal intensity using HCR v3.0 (blue, n=4 fish) or HCR-Immuno (red, n=4 fish). HCR-Immuno produced significantly higher net signal than HCR v3.0 at 50% and 100% laser power (p<0.0001 for both), whereas the difference at 10% laser power was not significant. **(E’)** Quantification of background fluorescence used to calculate the net signal shown in (**E**). Background fluorescence did not differ significantly at 10% laser power but was higher for HCR-Immuno at 50% (p < 0.01) and 100% (p < 0.001) laser power. **(F)** For HCR-Multi, first-round HCR amplifier hairpins (black) are conjugated to haptens (green circles) that are detected by a primary antibody (blue Y), followed by a secondary antibody (red Y) that is conjugated to an initiator (gray) for a second, orthogonal HCR amplifier system. This antibody-conjugated initiator triggers polymerization of the second-round HCR amplifier hairpins (black) which are conjugated to a different hapten (orange squares), which is detected by a second primary antibody (purple Y), followed by detection using a secondary antibody (light blue Y) that is conjugated to Alexa Fluor fluorophores (green stars). HCR-Multi produced a robust signal for *hcrt* (**H–H”**) compared to HCR v3.0 (**G–G”**) using 8 probes. (**I**) Quantification of net signal using HCR v3.0 (blue, n=4 fish) or HCR-Multi (red, n=4 fish). HCR-Multi produced significantly higher net signal than HCR v3.0 at all laser powers examined (p < 0.01 at 10% laser power, p < 0.0001 at 50% and 100% laser power). **(I’)** Quantification of background fluorescence used to calculate the net signal shown in (**I**). Background fluorescence did not differ significantly at 10% laser power but was higher with HCR-Multi at 50% (p < 0.01) and 100% (p < 0.0001) laser power. For (**C–D”**) and (**G–H”**), the same region was imaged at 10%, 50%, and 100% laser power while maintaining a fixed detector gain. All experiments used at least 4 fish. Mean ± SEM (arbitrary units, a.u.) are shown(**E, E’, I, I’**). Representative images shown are maximum intensity projections of z-stacks across the cell populations (**C–D”, G–H”**). Alexa Fluor 488 (**C,D**), Alexa Fluor 546 (**G**), and Alexa Fluor 568 (**H**) were used. Scale bars: 10 μm (**D”, H”**). Statistical significance was assessed using Two-way ANOVA with amplification method and laser power as factors, followed by Tukey’s multiple-comparisons test.

To further increase sensitivity while preserving spatial resolution, we developed HCR-Multi (Hybridization Chain Reaction with Multiple iterative amplification), which performs sequential rounds of HCR amplification using orthogonal amplifier systems and hapten-antibody pairs (**Fig. 3F**). Using *hcrt* as the target transcript in larval zebrafish, HCR-Multi produced markedly stronger fluorescence than HCR v3.0 across all laser powers tested (**Fig. 3G–H’’**). Quantification demonstrated an approximately 70-fold increase in net signal relative to HCR v3.0 (averaged across all laser powers tested) (**Fig. 3I**). As observed for HCR-Immuno, background fluorescence increased modestly with laser power (**Fig. 3I’**); however, the substantially larger increase in target signal resulted in a dramatic improvement in net signal. Together, these results demonstrate that HCR-Immuno and HCR-Multi provide complementary strategies for enhancing HCR sensitivity through immunodetection, with iterative amplification in HCR-Multi providing substantially greater signal enhancement than HCR-Immuno.

### HCR-Immuno and HCR-Multi enhance transcript detection in cleared mouse brain tissues

Having established that HCR-Immuno and HCR-Multi substantially improve mRNA detection sensitivity in whole-mount zebrafish larvae, we next evaluated their performance in thick PACT-cleared mouse brain sections using the same *YFP* transcript assay described for HCR-Cat (**Fig. 2B–E**). HCR-Immuno generated robust *YFP* transcript labeling throughout 600 μm-thick Thy1-YFP mouse brain sections using either FITC-or DIG-conjugated HCR amplifiers (**Fig. 4A**).

**Figure 4:**
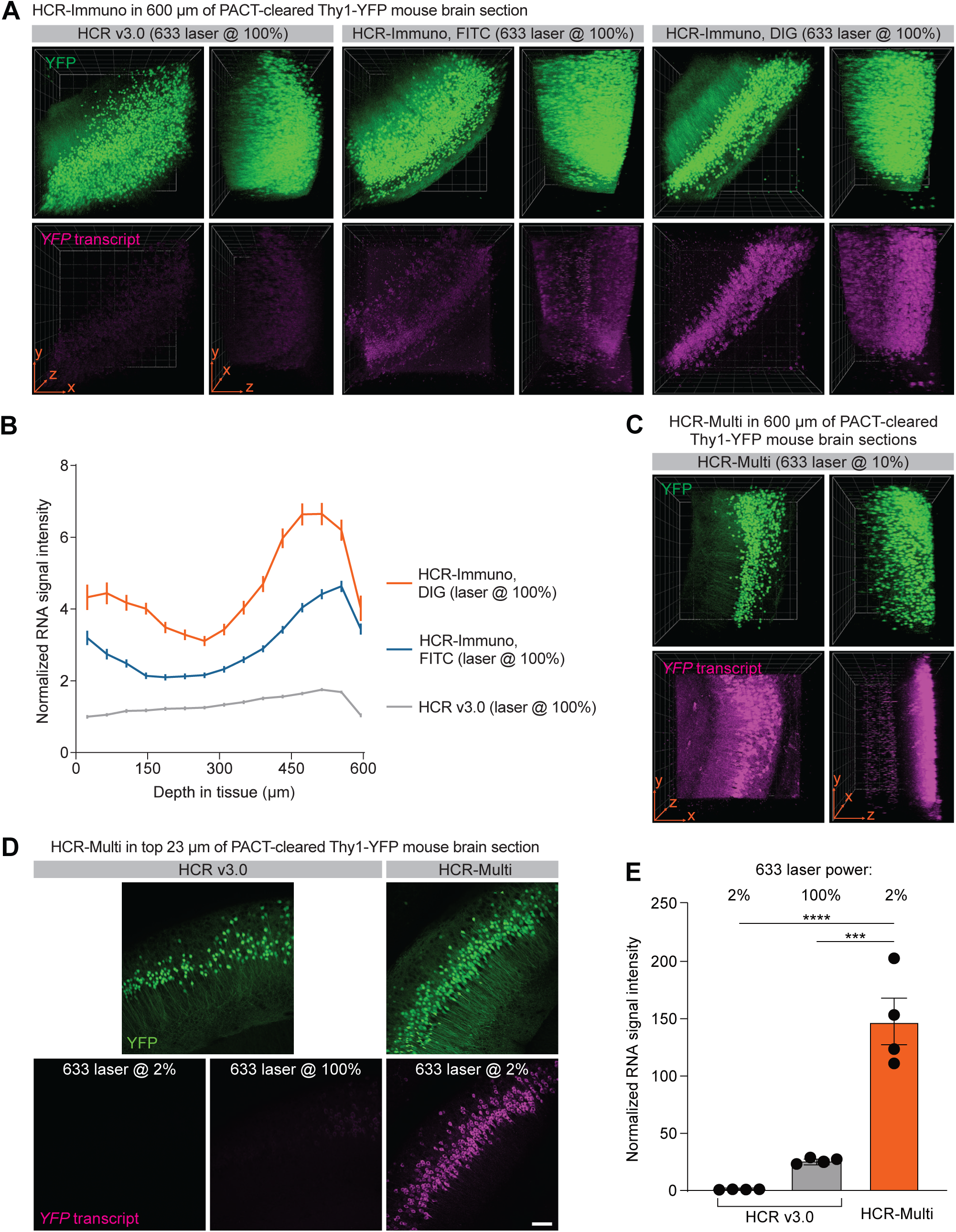
H**C**R**-Immuno and HCR-Multi in PACT-cleared thick mouse brain sections. (A)** Comparison of HCR v3.0 and HCR-Immuno for detection of *YFP* transcript (7 probes) in 600 μm-thick volumes of PACT-cleared Thy1-YFP mouse brain. Two HCR-Immuno detection strategies were tested: FITC-conjugated hairpins with a sheep anti-FITC primary antibody, and DIG-conjugated hairpins with a rabbit anti-DIG primary antibody. Both FITC- and DIG-based HCR-Immuno strategies enabled detection of *YFP* transcript throughout the tissue volume. No correction for imaging depth was applied. Representative images from 4 mice are shown. **(B)** Quantification of normalized *YFP* mRNA signal intensity as a function of tissue depth. RNA signal was quantified in YFP-positive cell bodies segmented using the YFP channel and normalized to the HCR v3.0 mRNA signal intensity at the tissue surface. HCR-Immuno using FITC yielded an approximately 3-fold increase (p < 0.05, at 23.8 μm, 64.6 μm, 431.8 μm, 472.6 μm, 513.4 μm, 554.2 μm, and 595 μm), whereas HCR-Immuno using DIG yielded an approximately 5-fold increase (p < 0.05, at all tissue depths) in RNA signal compared to HCR v3.0 when imaged at the same laser power and detector gain. Mean ± 99% confidence interval from 352-1516 cells per depth, pooled from 4 mice, is shown. Statistical significance was tested with a 2-way ANOVA with Dunnett’s multiple comparison test, treating the animal as the unit of replication. Complete statistical results, including adjusted p-values for pairwise comparisons, are provided in Table S1. **(C)** HCR-Multi in 600 μm-thick PACT-cleared Thy1-YFP mouse brain sections. HCR-Multi produced strong, but only superficial, *YFP* transcript staining. **(D)** Maximum intensity projections of the top 23 μm of PACT-cleared Thy1-YFP mouse brain stained using HCR v3.0 or HCR-Multi, showing robust transcript detection with HCR-Multi despite imaging at only 2% laser power. **(E)** Quantification of normalized RNA signal intensity at the tissue surface for HCR v3.0 and HCR-Multi. Mean ± SEM from 4 mice (132-431 cells per mouse) is shown. HCR-Multi yielded an approximately 150-fold increase in RNA signal compared to HCR v3.0 when imaged at the same 2% laser power and an approximately 6-fold increase compared to HCR v3.0 when imaged at 100% laser power. ***p = 0.0001, ****p < 0.0001. Statistical significance was tested with a One-way ANOVA with Tukey’s multiple comparison test, treating the animal as the unit of replication. Complete statistical results, including adjusted p-values for pairwise comparisons, are provided in Table S1. Scale bars: 160 μm (**A**, **C,** orange arrows in plane of image), 100 μm (**D**).

Quantification of RNA signal intensity throughout the tissue volume demonstrated that both detection strategies substantially outperformed HCR v3.0 across the full imaging depth, with the DIG-based implementation producing the highest signal (**Fig. 4B**). These results demonstrate that HCR-Immuno enables enhanced transcript detection throughout large, cleared tissue volumes and is compatible with multiple hapten-antibody detection chemistries.

In contrast, HCR-Multi produced strong transcript labeling that was largely restricted to the superficial region of the tissue (**Fig. 4C**), suggesting limited penetration of one or more amplification reagents during the sequential amplification workflow. To determine whether HCR-Multi nevertheless provides enhanced sensitivity within the labeled region, we quantified RNA signal within the most superficial 23 μm of the tissue volume. Despite imaging with only 2% laser power, HCR-Multi generated an approximately 150-fold increase in signal compared to HCR v3.0 imaged under the same conditions and remained approximately 6-fold brighter than HCR v3.0 at 100% laser power (**Fig. 4D,E**). These results demonstrate that HCR-Multi provides exceptional signal amplification in cleared tissue, but further optimization of reagent penetration is required for volumetric imaging of thick specimens.

### HCR-Cat provides the greatest signal enhancement among HCR-based amplification strategies

Having established that HCR-Cat, HCR-Immuno, and HCR-Multi each improve transcript detection relative to HCR v3.0, we next performed a direct comparison of all four amplification strategies in whole-mount larval zebrafish using the same *hcrt* probe set, identical imaging conditions, and the same range of laser powers. HCR-Cat generated the strongest fluorescence across the entire range of laser powers tested, followed by HCR-Multi, HCR-Immuno, and HCR v3.0 (**Fig. 5A–D”**). To facilitate visualization of the weaker signals produced by HCR-Multi, HCR-Immuno, and HCR v3.0, the corresponding images are also shown using an adjusted lookup table (LUT) to increase signal intensity while preserving the original image data (**Fig. 5B –D**).

**Figure 5:**
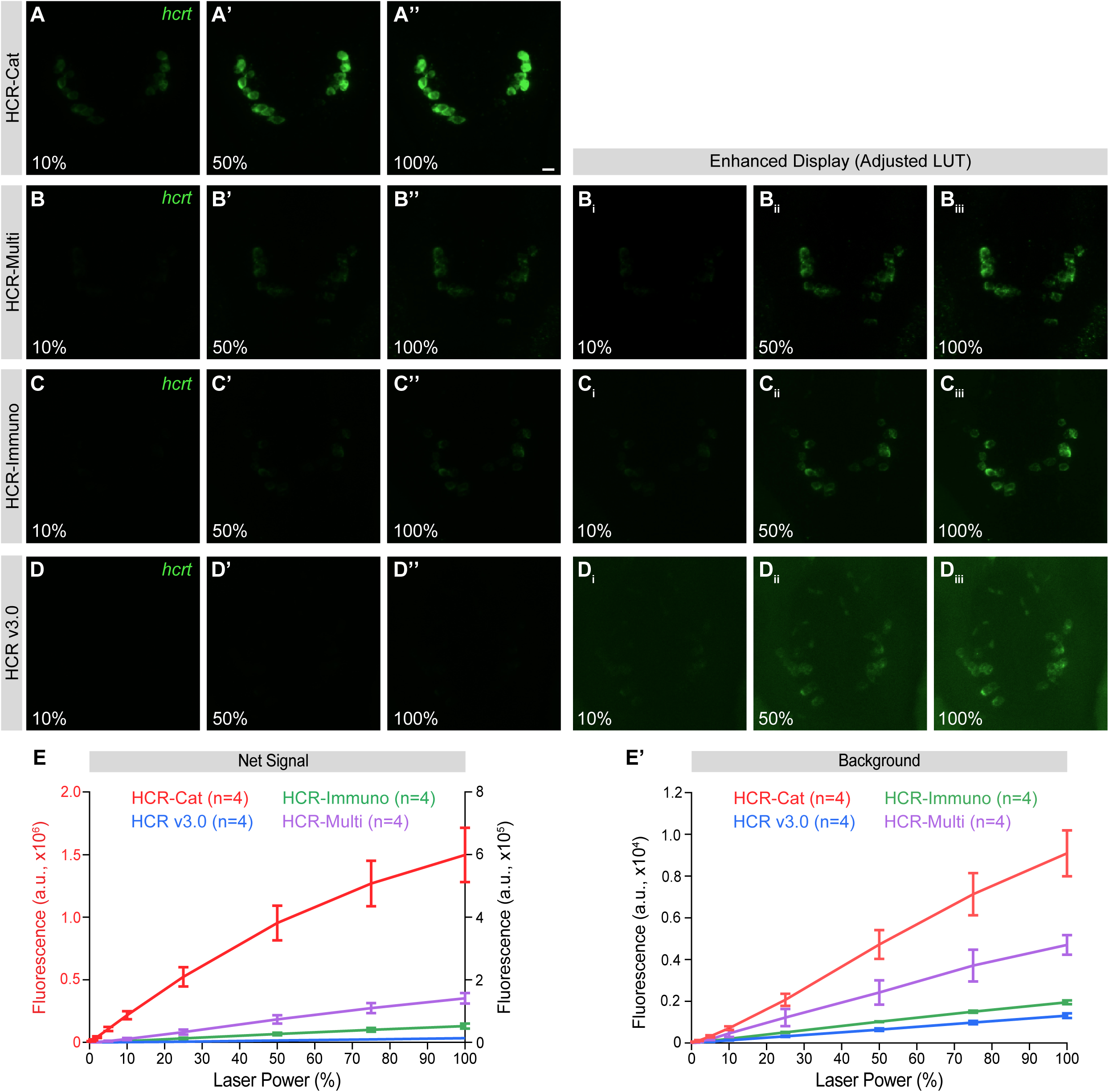
C**o**mparison **of fluorescence signal intensity generated by HCR-Cat, HCR-Multi, HCR-Immuno and HCR v3.0 in zebrafish larvae. (A-D)** Representative maximum intensity projections of z-stacks of *hcrt*-positive neurons in whole-mount zebrafish larvae detected using HCR-Cat (**A–A’’**), HCR-Multi (**B–B’’**), HCR-Immuno (**C–C’’**), or HCR v3.0 (**D–D’’**) using the same 8-probe *hcrt* probe set. Images were acquired at 10%, 50%, and 100% laser power while maintaining a fixed detector gain. HCR-Cat generated the strongest fluorescence signal across all laser powers, followed by HCR-Multi, HCR-Immuno, and HCR v3.0. Panels (**B_i_–B_iii_**), (**C_i_–C_iii_**) and (**D_i_–D_iii_**) show the HCR-Multi, HCR-Immuno, and HCR v3.0 images using an increased LUT to facilitate visualization of the weaker fluorescence signals. Alexa Fluor 488 (**B–D**) and TSA-Flu (**A**) were used. Scale bar: 10 μm (**A”**). **(E)** Quantification of net fluorescence signal as a function of laser power for HCR-Cat (red), HCR-Multi (purple), HCR-Immuno (green), and HCR-v3.0 (blue). HCR-Cat is quantified using the left y-axis. The other three conditions are quantified using the right y-axis. All experiments used at least 4 fish. **(E’)** Quantification of background fluorescence used to calculate the net signal shown in (**E**). For each sample, the region of interest was imaged by gradually increasing the laser power from 0% to 100% while maintaining a fixed detector gain. Mean ± SEM (arbitrary units, a.u.) are shown (**E, E’**). Statistical significance was assessed using two-way ANOVA with amplification method and laser power as factors, followed by Tukey’s multiple-comparisons test. Pairwise comparisons between amplification methods at each matched laser power were performed; complete statistical results, including adjusted P values, are provided in Table S1.

Quantification confirmed these qualitative observations (**Fig. 5E**). HCR-Cat produced the highest net fluorescence signal at every laser power examined, followed by HCR-Multi, HCR-Immuno, and HCR v3.0. Although background fluorescence increased with laser power for all four methods (**Fig. 5E’**), the increase in target signal was substantially greater, resulting in the highest net signal for HCR-Cat across the full range of imaging conditions tested. HCR-Multi offers substantial signal enhancement while preserving spatial resolution, whereas HCR-Immuno provides increased sensitivity compared to HCR v3.0 through a simpler antibody-based workflow compared to HCR-Multi while maintaining spatial resolution. Together, these results establish clear performance trade-offs that can guide selection of the appropriate amplification strategy for different experimental applications.

### Enhanced transcript detection is maintained under single-probe pair conditions

Many biologically important RNA targets accommodate only a limited number of probe pairs, which can be challenging to detect using HCR v3.0. Having established the relative performance of the four amplification strategies under conventional multi-probe conditions (**Fig. 5**), we next asked whether these relative advantages are maintained when only a single split-probe pair is used. To do so, we directly compared HCR-Cat, HCR-Immuno, HCR-Multi, and HCR v3.0 using one randomly selected split-probe pair targeting *dbh*, without prior optimization under identical imaging conditions (**Fig. 6A–D”**). Using a single probe pair substantially reduced fluorescence intensity for all four methods. Nevertheless, HCR-Cat generated the strongest fluorescence signal across the full range of laser powers tested, followed by HCR-Multi, HCR-Immuno, and HCR v3.0. To facilitate visualization of the weaker fluorescence generated by HCR-Multi, HCR-Immuno, and HCR v3.0, the corresponding images are also shown using an adjusted LUT to increase signal intensity while preserving the original image data (**Fig. 6B – D**).

**Figure 6:**
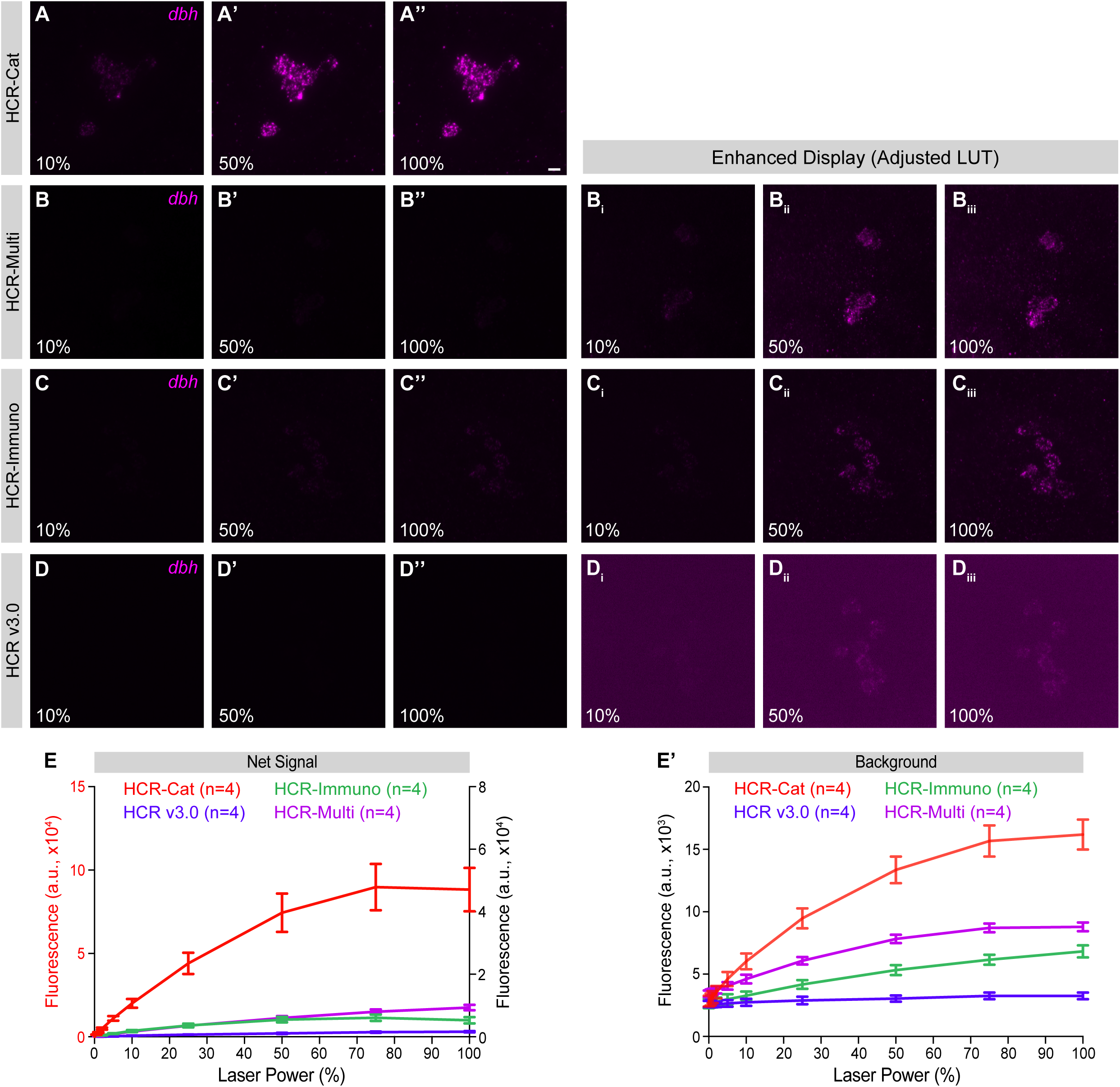
C**o**mparison **of single-probe detection sensitivity of HCR-Cat, HCR-Multi, HCR-Immuno and HCR v3.0 in zebrafish larvae. (A–D)** Representative maximum intensity projections of z-stacks across *dbh*-positive neurons in the locus coeruleus in whole-mount zebrafish larvae detected using HCR-Cat (**A–A’’**), HCR-Multi (**B–B’’**), HCR-Immuno (**C–C’’**), or HCR v3.0 (**D–D’’**) using a single *dbh* split-probe pair. Images were acquired at 10%, 50%, and 100% laser power while maintaining a fixed detector gain. HCR-Cat generated the strongest fluorescence signal across all laser powers, followed by HCR-Multi, HCR-Immuno, and HCR v3.0. Panels (**B_i_–B_iii_**), (**C_i_–C_iii_**) and (**D_i_–D_iii_**) show the HCR-Multi, HCR-Immuno, and HCR v3.0 images using an increased LUT to facilitate visualization of the weaker fluorescence signals. Alexa Fluor 633 (**B–D**) and TSA-Cy5 (**A**) were used. Scale bar: 10 μm (**A”**). **(E)** Quantification of net fluorescence signal as a function of laser power for HCR-Cat (red), HCR-Multi (purple), HCR-Immuno (green), and HCR-v3.0 (blue). HCR-Cat is quantified using the left y-axis. The other three conditions are quantified using the right y-axis. All experiments used at least 4 fish. **(E’)** Quantification of background fluorescence used to calculate the net signal shown in (**E**). For each sample, the region of interest was imaged by gradually increasing the laser power from 0% to 100% while maintaining a fixed gain. Mean ± SEM (arbitrary units, a.u.) are shown (**E, E’**). Statistical significance was assessed using two-way ANOVA with amplification method and laser power as factors, followed by Tukey’s multiple-comparisons test. Pairwise comparisons between amplification methods at each matched laser power were performed; complete statistical results, including adjusted P values, are provided in Table S1.

Quantification confirmed that the relative performance of the four amplification strategies was preserved under single-probe conditions (**Fig. 6E**). HCR-Cat produced the highest net fluorescence signal at every laser power examined, whereas HCR-Multi and HCR-Immuno generated lower signal levels that were still higher than HCR v3.0. Although background fluorescence increased with laser power for all four methods (**Fig. 6E’**), the increase in target signal remained significantly greater for each of the three amplification strategies compared to HCR v3.0, resulting in markedly improved net signal across all imaging conditions. Together, these findings demonstrate that HCR-Cat, HCR-Immuno, and HCR-Multi provide enhanced detection sensitivity even when only a single split-probe pair is used, supporting their application to RNA targets for which probe design is constrained, including short transcripts and other challenging RNA species.

### Split-probe specificity is maintained in HCR-Cat, HCR-Immuno, and HCR-Multi

To confirm that the enhanced sensitivity of HCR-Cat, HCR-Immuno, and HCR-Multi does not compromise the split-probe specificity of HCR v3.0, we next performed each amplification strategy using only one member of each split-probe pair (referred to here as the odd- or even-half-probe set), which will not generate a complete HCR initiator, and thus should not allow binding of a hairpin, for detection of *hcrt* in whole-mount zebrafish larvae (**Fig. S6**). Consistent with HCR v3.0, none of the three amplification strategies produced specific labeling of *hcrt-* expressing neurons when only the odd- or even-half-probe set was used. Although occasional bright puncta were observed outside the expected expression domain, these did not exhibit the characteristic anatomical distribution of *hcrt*-expressing neurons and are likely due to low-level non-specific binding of antibodies to the surface of the fish. These results demonstrate that signal generation by HCR-Cat, HCR-Immuno, and HCR-Multi, like HCR v3.0, remains dependent on complete split-probe assembly, confirming that the enhanced sensitivity of the three amplification strategies is achieved without compromising split-probe-dependent detection specificity.

### HCR-Immuno preserves transcript-resolution detection following short amplification times

Because HCR-Immuno enhances fluorescence signal while in principle preserving transcript localization, we reasoned that it could be particularly useful for applications requiring transcript-resolution detection. HCR v3.0 typically achieves the most robust transcript-resolution detection using relatively large probe sets, often comprising dozens of split-probe pairs. Consequently, application to short RNA targets that accommodate only a limited number of probe pairs remains challenging. However, the antibody-dependent signal enhancement of HCR-Immuno could potentially alter puncta morphology and compromise transcript-resolution detection. To determine if this is the case, we used the complete 20-pair *dbh* split-probe set, but divided the probes into two independent subsets (10 probe pairs per subset). In contrast to the split-probe specificity experiment described above, each subset contained complete split-probe pairs, with both probes required to form the HCR initiator. Each subset was detected using a different HCR amplifier system and fluorescence channel. Colocalization of puncta detected in the two channels provides a well-established criterion for identifying bona fide single-transcript signals (Choi et al., 2018). HCR v3.0 and HCR-Immuno were performed using HCR amplification times of 15, 30, or 60 minutes (**Fig. 7A-B”**). Both methods produced well-resolved puncta following 15 minutes of amplification, whereas progressively longer amplification times resulted in increased puncta clustering and larger fluorescent structures, indicating that shorter amplification times better preserve transcript-resolution detection.

**Figure 7:**
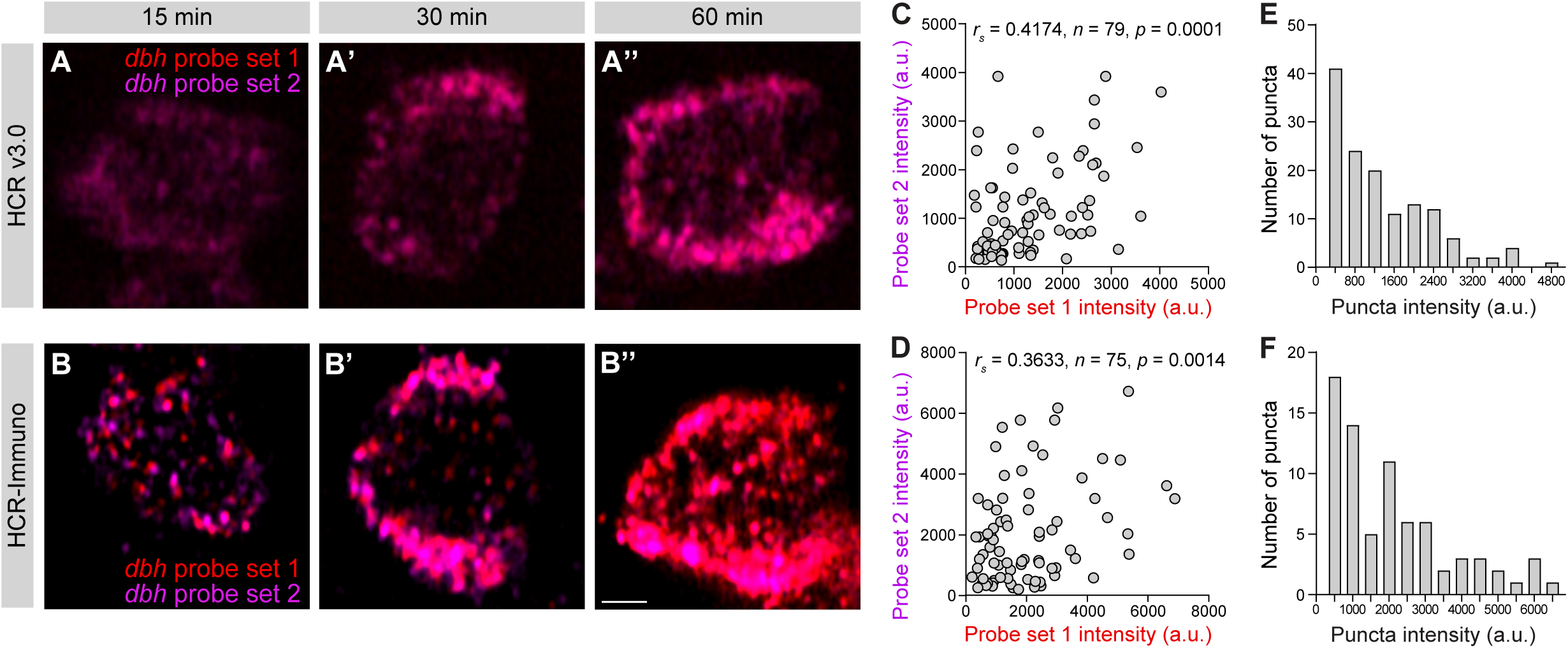
H**C**R**-Immuno preserves puncta morphology following short amplification times in larval zebrafish. (A–A’’)** Representative images of single *dbh*-positive neurons in larval zebrafish detected using HCR v3.0 using two different *dbh* probe sets of 10 probes each (probe subset 1 = red, probe subset 2 = purple) with 15 min (**A**), 30 min (**A’**), or 60 min (**A’’**) amplification. **(B–B’’)** Corresponding HCR-Immuno images acquired using the same *dbh* probe subsets following 15 min (**B**), 30 min (**B’**), or 60 min (**B’’**) amplification. Single optical sections are shown. Puncta were well resolved after 15 min amplification but not after longer amplification times for both methods. Alexa Fluor 546 and 633 were used for probe subsets 1 and 2, respectively for HCR v3.0 (**A–A”**). Alexa Fluor 568 and Alexa Fluor 633 were used for probe subsets 1 and 2, respectively, in HCR-Immuno (**B–B”**). Scale bar: 2.5 μm **(B’’)**. **(C–D)** Scatter plots show fluorescence intensity of matched puncta for probe subsets 1 and 2 using HCR v3.0 (**C**) and HCR-Immuno (**D**). Matched puncta showed a positive and statistically significant correlation by Spearman correlation analysis (*r*) for both HCR v3.0 and HCR-Immuno. n = number of puncta analyzed. **(E–F)** Frequency distribution of puncta fluorescence intensity for HCR v3.0 (**E**) and HCR-Immuno (**F**). Both distributions were unimodal, consistent with a predominant population of similarly amplified transcript-associated puncta. Unimodality was confirmed by Hartigan’s dip test (HCR v3.0: dip = 0.0190, p = 0.990; HCR-Immuno: dip = 0.0303, p = 0.837). Puncta fluorescence intensity is shown in arbitrary units (a.u.). Quantification was performed using the 15 min amplification dataset.

To quantitatively compare transcript-resolution detection, we analyzed the 15-minute amplification condition by matching puncta detected by both of the 2 probe subsets using nearest-neighbor centroid analysis. For both HCR v3.0 and HCR-Immuno, significant correlations were observed between matched puncta in the two channels for fluorescence intensity (**Fig. 7C,D**) and puncta area (**Fig. S7A,B**). Although the degree of correspondence between the two probe subsets was lower than previously reported for optimized HCR v3.0 probe sets (Choi et al., 2018), detection by independent probe subsets provides an estimate of probe-detection efficiency rather than, by itself, establishing that individual puncta represent single transcripts. Importantly, the fluorescence intensity distributions of detected puncta were unimodal for both HCR v3.0 and HCR-Immuno (**Fig. 7E,F**), consistent with a predominant population of similarly amplified RNA molecules rather than distinct populations representing aggregated transcripts. Puncta size distributions were likewise unimodal and comparable between the two methods (**Fig. S7C,D**). Together with the discrete, well-resolved punctate signals observed after 15 minutes of amplification, these quantitative measurements are consistent with transcript-resolution detection by HCR-Immuno and demonstrate that antibody-mediated amplification can enhance fluorescence while preserving the punctate characteristics of HCR v3.0.

### Compatibility with diverse detection modalities and HCR amplifier chemistries

To expand the versatility of the amplification strategies described here, we evaluated whether HCR-Cat is compatible with alkaline phosphatase (AP)-mediated chromogenic detection, which remains widely used for *in situ* hybridization in developmental biology, histology, and pathology. Using *dbh* as the target transcript, we tested two commonly used AP substrates, Vector Red and nitro blue tetrazolium/5-bromo-4-chloro-3-indolyl-phosphate (NBT/BCIP), in the presence or absence of HCR probes (**Fig. S8**). Robust chromogenic labeling of *dbh*-expressing neurons was observed with both substrates in probe-containing samples, whereas no signal was detected in the absence of probes. These results demonstrate that HCR-Cat is compatible with both fluorescent and brightfield AP-based chromogenic detection, extending its applicability to conventional chromogenic *in situ* hybridization workflows without compromising probe-dependent specificity.

To determine whether antibody-mediated amplification requires hapten-conjugated HCR amplifier hairpins, we next evaluated HCR-Immuno using Alexa Fluor 488-conjugated HCR amplifier hairpins together with an anti-Alexa Fluor 488 antibody (**Fig. S9**). Under identical imaging conditions, HCR-Immuno produced substantially greater fluorescence than HCR v3.0 across all laser powers tested. These results demonstrate that HCR-Immuno is compatible with Alexa Fluor 488-conjugated HCR amplifier hairpins, providing an alternative to the hapten-conjugated amplifier hairpins used elsewhere in this study and broadening the range of HCR amplifier reagents that can be used for antibody-mediated signal enhancement.

## Discussion

Spatial transcriptomic methods often face a fundamental trade-off between detection sensitivity and transcript-resolution imaging. Here, we introduce three complementary HCR-based amplification strategies that substantially extend the capabilities of HCR v3.0 while retaining its split-probe specificity. HCR-Cat provides the greatest increase in detection sensitivity through catalytic reporter deposition, enabling robust detection of transcripts that are expressed at low levels, under conditions where only limited probe occupancy is available, and in highly autofluorescent or thick samples. HCR-Immuno increases fluorescence while preserving transcript-resolution detection through antibody-mediated signal amplification, whereas HCR-Multi further enhances signal through iterative rounds of HCR and immunodetection. Together, these approaches provide a flexible signal amplification toolkit that allows sensitivity, spatial resolution, and workflow complexity to be balanced according to the requirements of individual biological applications.

A direct comparison of all four amplification strategies under identical experimental conditions established a clear hierarchy of fluorescence enhancement, with HCR-Cat providing the greatest signal amplification, followed by HCR-Multi, HCR-Immuno, and HCR v3.0. Importantly, this comparison demonstrates that the three amplification strategies occupy distinct positions along the sensitivity–resolution spectrum rather than representing competing alternatives. HCR-Cat is ideally suited for applications where maximal sensitivity is required, particularly in highly autofluorescent or optically challenging tissues. HCR-Immuno preserves transcript-resolution detection while providing increased fluorescence signal through a simple antibody-based workflow, making it well suited for multiple transcript imaging where increased fluorescence and preservation of spatially resolved puncta are desirable. HCR-Multi provides an intermediate strategy for experiments requiring greater sensitivity than HCR-Immuno, while avoiding the catalytic reporter deposition of HCR-Cat. Rather than replacing HCR v3.0, these approaches broaden the experimental situations in which HCR-based transcript detection can be successfully applied. The approaches also differ in their reagent requirements, workflow complexity, and practical costs. HCR-Immuno requires only an additional antibody-detection step following HCR amplification, whereas HCR-Multi requires additional amplifier reagents and sequential rounds of antibody labeling and HCR amplification, increasing both reagent use and hands-on time. HCR-Cat requires enzyme-conjugated antibodies and reporter-deposition reagents but provides the greatest signal enhancement without iterative rounds of HCR amplification. Thus, selection of an amplification strategy will depend not only on the required sensitivity and spatial resolution, but also on reagent availability, experimental scale, workflow time, and cost.

An important practical implication of our findings is the compatibility of HCR-Immuno with Alexa Fluor 488-conjugated HCR amplifier hairpins detected using an anti-Alexa Fluor 488 antibody. This expands the range of amplifier reagents that can be used for antibody-mediated signal enhancement beyond those used in our standard HCR-Immuno workflow. Because HCR-Cat and HCR-Multi similarly rely on antibody recognition of amplifier-associated labels, fluorophore-conjugated HCR amplifier hairpins should in principle also be compatible with these approaches when suitable fluorophore-specific antibodies are available. This flexibility could facilitate adaptation of these amplification strategies to existing HCR workflows and reagent sets. Because these amplification strategies operate downstream of probe hybridization, existing HCR probe libraries can in principle be used without redesign. However, their practical multiplexing capacity is constrained by the availability of mutually compatible amplifier-associated labels and corresponding label-specific antibodies. Combining orthogonal hapten- and fluorophore-based detection strategies may provide a route to increased multiplexing, but higher-order multiplexing was not directly evaluated in this study.

One important advance of this study is the demonstration that enhanced sensitivity is maintained under highly constrained probe conditions. Many biologically important RNA targets accommodate only a limited number of split-probe pairs, making sensitive detection challenging. We show that HCR-Cat, HCR-Immuno, and HCR-Multi all produce substantial signal enhancement even when only a single split-probe pair is used. Although we did not directly examine short RNA species such as microRNAs, our findings suggest that the methods described here could enhance detection of short or otherwise sequence-constrained RNA targets that can accommodate at least one split-probe pair. Mature miRNAs, however, are typically only ∼20–24 nucleotides long and may not accommodate the standard split-probe architecture used in this study. Application of these amplification strategies to mature miRNAs would therefore require additional methodological development and experimental validation.

Because increased amplification can potentially compromise transcript-resolution detection, we evaluated whether HCR-Immuno preserves the spatial precision of HCR v3.0. Using an independent probe-subset colocalization strategy to estimate detection efficiency, together with quantitative analyses of puncta morphology and fluorescence intensity, we found that HCR-Immuno maintained transcript-resolution imaging following short amplification times while providing substantially greater fluorescence signal than HCR v3.0. Importantly, correspondence between independently detected probe subsets provides an estimate of detection efficiency rather than, by itself, establishing single-transcript detection (Choi et al., 2018; Ji and van Oudenaarden, 2012). Instead, the preservation of discrete, well-resolved puncta, together with fluorescence intensity distributions showing no significant departure from unimodality, provides complementary evidence consistent with a predominant population of individual transcript-associated signals (Choi et al., 2018; Ji and van Oudenaarden, 2012). Similar puncta size distributions, which likewise showed no significant departure from unimodality, further indicate that antibody-mediated amplification does not alter puncta morphology. These findings indicate that antibody-mediated signal enhancement can be incorporated into HCR without compromising transcript-resolution information. However, we have not systematically determined how amplification scales with transcript abundance, probe-set size, or amplification duration, and therefore cannot conclude whether signal amplification is linear or follows another quantitative relationship under these conditions. Systematic characterization of amplification scaling across different targets, probe-set sizes, and experimental conditions will be important for applications requiring quantitative comparison or higher-order multiplexing.

Several approaches have previously been developed to improve HCR sensitivity. USeqFISH (Jang et al., 2023) combines rolling-circle amplification with HCR and sequential imaging to achieve high sensitivity with relatively few probes while enabling large-scale multiplexing. However, this approach requires enzymatic amplification, tissue clearing, and a substantially more complex workflow. Alternative strategies, including shortened hairpins (Tsuneoka and Funato, 2020), branched hairpins (Xu et al., 2019), and additional hairpin purification (Leino and Söderberg, 2023) improve HCR polymerization efficiency but require modified hairpin designs or specialized reagent preparation. In contrast, our amplification strategies are fully compatible with existing HCR v3.0 probe design principles and workflows, requiring only modification of the amplification stage. HCR-Immuno and HCR-Multi retain the enzyme-free simplicity of HCR amplification, whereas HCR-Cat uses HRP-conjugated antibodies to achieve catalytic signal deposition without the additional enzymatic amplification steps required by rolling-circle amplification-based approaches.

Although HCR-Immuno performed well in whole-mount zebrafish and thin mouse brain sections, antibody penetration is expected to become increasingly limiting in large tissue volumes. Indeed, HCR-Multi showed reduced penetration in 600-µm PACT-cleared mouse cortex, likely reflecting the cumulative diffusion barriers imposed by repeated rounds of antibody labeling. These limitations may be addressed through longer incubations, optimized tissue-clearing protocols, or smaller affinity reagents such as Fab or F(ab′)_₂_ fragments (Guerin, 2023) or nanobodies (Yamauchi et al., 2025), which should improve penetration while preserving the modular nature of the amplification strategy.

The amplification strategies described here are compatible with both hydrogel-based (PACT) and solvent-based (iDISCO+) tissue-clearing approaches, demonstrating their applicability across chemically distinct clearing methods. More generally, these approaches should be compatible with clearing protocols that preserve RNA integrity, HCR amplification products, and antibody accessibility. As with conventional immunostaining, optimization of tissue permeability, reagent penetration, antibody incubation times, and refractive index matching may be required depending on tissue type, sample size, and the clearing method employed. These considerations are expected to be particularly important for HCR-Immuno and HCR-Multi, where antibody diffusion contributes directly to amplification efficiency.

An additional strength of the approaches described here is their flexibility. We demonstrate compatibility with multiple hapten-antibody systems, alkaline phosphatase-mediated chromogenic detection, and fluorophore-conjugated HCR amplifier hairpins detected using fluorophore-specific antibodies. Because all three amplification strategies operate downstream of probe hybridization, they are compatible with existing HCR probe libraries, allowing users to increase detection sensitivity by modifying only the post-hybridization detection chemistry. The modularity of initiator-conjugated secondary antibodies provides a flexible framework for future multiplex immunolabeling applications in which independently addressable HCR amplification can be assigned to different antibody targets. Beyond RNA detection, the same amplification framework should be readily adaptable to conventional immunohistochemistry by coupling initiator-conjugated secondary antibodies to protein-labeling workflows. Because HCR-Multi iteratively amplifies antibody-derived HCR signals, similar strategies could provide substantial enhancement of protein detection while preserving the modular architecture of HCR-based amplification.

In summary, HCR-Cat, HCR-Immuno, and HCR-Multi substantially expand the capabilities of HCR-based transcript detection by providing complementary solutions for experiments requiring increased sensitivity while preserving the specificity of split-initiator HCR. Together, these methods extend HCR to challenging biological applications that were previously difficult or impractical using HCR v3.0, while maintaining compatibility with existing HCR workflows and providing a flexible framework for future methodological development.

## Materials and Methods

### Zebrafish

#### Zebrafish husbandry

All zebrafish experiments were performed in accordance with protocols approved by the Institutional Animal Care and Use Committee (IACUC; protocol 1836) and the Office of Laboratory Animal Resources at the California Institute of Technology. Unless otherwise indicated, experiments were performed using 5-7 days post fertilization (dpf) larval TLAB zebrafish, generated by crossing AB and TL strains obtained from the Zebrafish International Resource Center. Embryonic and larval zebrafish were raised at 28.5°C in E3 embryo medium (5 mM NaCl, 0.17 mM KCl, 0.33 mM CaCl_2_, 0.33 mM MgSO_4_, pH 7.4) in petri dishes.

#### Common HCR workflow for zebrafish samples

Unless otherwise indicated, all HCR-based staining methods followed the common workflow described below. HCR probes, amplifier hairpins (B1–B5) (Choi et al., 2018), buffers, and initiator-labeled secondary antibodies were purchased from Molecular Technologies and Molecular Instruments. Alexa Fluor-conjugated HCR amplifier hairpins used as controls and in fluorophore-specific antibody experiments were obtained from Molecular Technologies. The zebrafish probe sets used in this study were commercially designed and are available for purchase from Molecular Technologies. As alternatives to commercially obtained reagents,

HCR probes can be designed using published scripts (Kroll et al., 2025; Kuehn et al., 2022), amplifier hairpins synthesized according to published methods (Choi et al., 2018; Leino and Söderberg, 2023), and buffers can be prepared using published recipes (Choi et al., 2018; Ćorić et al., 2023), and initiator-conjugated antibodies for HCR-Multi can be prepared using previously described antibody–oligonucleotide conjugation methods (Schwarzkopf et al., 2021). Throughout this manuscript, a “split-probe pair” refers to two oligonucleotide probes that bind adjacent sequences on the target RNA to generate a complete HCR initiator, whereas a “probe set” refers to multiple split-probe pairs targeting different regions of the same RNA (Choi et al., 2018). Staining was performed according to a previously published protocol (Shainer et al., 2023) with the modifications described below.

Typically, 10-12 zebrafish larvae were processed in 1.5-mL microcentrifuge tubes, using 500 μL of solution for each incubation and wash step. Unless otherwise indicated, incubations and washes were performed with gentle shaking. Larvae were anesthetized in 1.5 mM tricaine (Sigma-Aldrich, E10521) and fixed overnight at 4°C in ice-cold 4% paraformaldehyde (PFA) (Electron Microscopy Sciences, #15714-S) prepared in 1× DPBST (1× Dulbecco’s phosphate-buffered saline without divalent cations containing 0.1% Tween-20). Samples were then washed three times for 5 min in DPBST and bleached with 1.5% hydrogen peroxide, 1% potassium hydroxide, and 0.1% Tween-20 for 30 min at room temperature while monitoring pigment loss. Following five washes in DPBST, larvae were incubated in DPBSTx (1× DPBS containing 0.25% Triton X-100) for 1 h at room temperature to increase sample permeability. Samples were then dehydrated in chilled 100% methanol at −20°C for 10 min, followed by rehydration through 50% and 25% methanol in DPBSTx for 5 min each at room temperature, and then five additional washes in DPBSTx.

Samples were then equilibrated in 10% hybridization buffer in 5× sodium chloride sodium citrate + 0.1% Tween 20 (SSCT) for at least 15 min at 37°C, followed by pre-hybridization in 100% hybridization buffer for 1–2 h. The pre-hybridization buffer was then replaced with pre-warmed probe solution (100% hybridization buffer with 16-20 nM probes for weakly expressed transcripts or 2-4 nM probes for highly expressed transcripts) and incubated overnight at 37°C. For weakly expressed targets, hybridization for 36–60 h generally improved fluorescence signal. Following hybridization, samples were washed four times for 15 min each in pre-warmed probe wash buffer at 37°C, followed by two 5-min washes in 5× SSCT at room temperature.

For HCR amplification, samples were equilibrated in amplification buffer for 1–2 h at room temperature. Amplifier hairpins (30 pmol each) were snap-cooled individually by heating to 95°C for 90 s, followed by passive cooling to room temperature in the dark for at least 30 min. Hairpins were then combined in amplification buffer and incubated with the samples for 12–16 h at room temperature in the dark without shaking. Excess hairpins were removed by four 15-min washes in 5× SSCT. At this point, the protocol diverged according to the signal detection strategy described below.

#### HCR v3.0 for larval zebrafish samples

After the common HCR workflow, samples were washed once in 1× DPBS for 10 min before equilibration through 10%, 25%, 50%, and 80% VECTASHIELD PLUS (Vector Laboratories, H-1900-10) in 1× DPBS for at least 15 min at each concentration. Samples were imaged in 80% VECTASHIELD PLUS.

#### HCR-Cat for larval zebrafish samples

After the common HCR workflow, samples were washed three times in DPBSTx and then blocked for 1 h in Roche Blocking Solution (Roche, 11585762001). FITC-conjugated amplifier hairpins were detected using anti-Fluorescein-horseradish peroxidase (anti-Flu-HRP) Fab fragments (Roche, 11426346910; 1:400) overnight at 4°C in blocking solution, followed by five 15-min washes in DPBSTx. Signal was developed for 30 min using TSA PLUS Fluorescein (Akoya Biosciences, NEL741001KT) diluted 1:200 for weakly expressed targets or 1:300 for highly expressed targets in TSA amplification diluent. Following signal development, samples were washed five times for 5 min each in DPBSTx, followed by one wash in 1× DPBS for 10 min. Samples were then incubated overnight in DPBSTx at 4°C to reduce background fluorescence before equilibration in VECTASHIELD PLUS, as described above.

For DIG-based HCR-Cat, FITC-conjugated amplifier hairpins and anti-Flu-HRP Fab fragments were replaced with DIG-conjugated amplifier hairpins, anti-Digoxigenin-HRP (anti-DIG-HRP) Fab fragments (Roche, 11207733910, 1:400), and TSA PLUS Cyanine 5 (Akoya Biosciences, NEL745001KT), while all remaining steps were unchanged. Signal was developed for 30 min using TSA PLUS Cyanine 5 diluted 1:200 for weakly expressed targets or 1:300 for highly expressed targets in TSA amplification diluent.

For dual-target HCR-Cat, the first target was detected using FITC-conjugated HCR amplifier hairpins, anti-Flu-HRP, and TSA PLUS Fluorescein as described above. Residual HRP activity following the first TSA reaction was inactivated using 0.02 N HCl for 30 min (Liu et al., 2006), after which a second round of blocking, anti-DIG-HRP incubation, and TSA PLUS Cyanine 5 development was performed using the same procedure.

Alternatively, FITC-conjugated amplifier hairpins were detected using anti-Fluorescein-alkaline phosphatase (anti-Flu-AP) Fab fragments (Roche, 11426338910; 1:2000) overnight at 4°C. Following five 15-min washes in DPBSTx, signal was visualized using either ImmPACT Vector Red Alkaline Phosphatase Substrate (Vector Laboratories, SK-5105) according to the manufacturer’s instructions or NBT/BCIP (Roche, 11383213001 and 11383221001). For NBT/BCIP detection, samples were equilibrated three times for 5 min each in NTMT buffer (100 mM Tris-HCl, pH 9.5, 50 mM MgCl_₂_, 100 mM NaCl, and 0.1% Tween-20), followed by incubation in NBT/BCIP staining solution (45 μL NBT and 35 μL BCIP in 10 mL NTMT buffer) at room temperature in the dark. The staining reaction was terminated by five rapid washes in DPBSTx, after which samples were post-fixed in 4% PFA before equilibration in VECTASHIELD PLUS, as described above.

#### HCR-Immuno for larval zebrafish samples

After the common HCR workflow, samples were washed three times in DPBSTx and blocked for 1 h in 2% normal goat serum and 2% DMSO in DPBSTx. DIG-conjugated amplifier hairpins were detected using rabbit anti-DIG antibody (Invitrogen, 700772; 1:100) overnight at 4°C, followed by five washes in DPBSTx and incubation with an Alexa Fluor-conjugated goat anti-rabbit secondary antibody (Thermo Fisher, A-11008, 1:500) overnight at 4°C. Samples were washed and equilibrated in VECTASHIELD PLUS as described above. For transcript-resolution experiments, HCR amplification was performed for 15, 30, or 60 min at room temperature before immunodetection. The HCR-Immuno experiments shown in Figure 3 were performed using ethylene carbonate-based hybridization buffers (all % v/v): Hybridization buffer: 2× SSC, 10% ethylene carbonate (Sigma-Aldrich, E26258), 10% dextran sulfate (Sigma-Aldrich, #3730), and 0.1% Tween-20; Stringent wash buffer: 2× SSC, 30% ethylene carbonate and 0.1% Tween-20; Wash buffer: 2× SSC and 0.1% Tween-20; Amplification buffer: 2× SSC, 10% ethylene carbonate and 0.1% Tween-20. These buffers provided improved signal-to-noise compared with conventional formamide-based buffers while avoiding the hazards associated with formamide (Matthiesen and Hansen, 2012). For experiments using fluorophore-conjugated HCR amplifier hairpins, Alexa Fluor 488-conjugated hairpins were detected using an anti-Alexa Fluor 488 primary antibody (Thermo Fisher, A-11094, 1:100) followed by an Alexa Fluor 488-conjugated goat anti-rabbit IgG (H+L), cross-adsorbed secondary antibody (Thermo Fisher, A-11008, 1:500) using the same blocking, incubation, and washing conditions described above. All other HCR-Immuno experiments were done with the standard buffers described above.

#### HCR-Multi for larval zebrafish samples

After the common HCR workflow, DIG-conjugated B1 amplifier hairpins were detected using rabbit anti-DIG antibody (Invitrogen, 700772; 1:100), followed by an initiator-conjugated donkey anti-rabbit secondary antibody carrying the B2 initiator (1:500). Following antibody incubation and washing, a second round of HCR amplification was performed using FITC-conjugated B2 hairpins, followed by detection with mouse anti-FITC antibody (Invitrogen, 31242; 1:500) and an Alexa Fluor 568-conjugated goat anti-mouse IgG (H+L), cross-adsorbed secondary antibody (Thermo Fisher, A-11004, 1:500). Samples were then washed and mounted in VECTASHIELD PLUS as described above.

#### Imaging, quantification, and data analysis for larval zebrafish samples

Samples were mounted in 80% VECTASHIELD PLUS and imaged using a Zeiss LSM 880 confocal microscope. Individual fish were mounted dorsal side up within a well formed by 5–6 reinforcement labels on a glass slide, with the viscosity of the mounting medium maintaining sample orientation. Low-magnification images were acquired using a 20×/1.0 NA water-dipping objective (W Plan-Apochromat 20×/1.0 DIC CG = 0.17 M27, 75 mm) with the Zeiss tiling function and stitched using the Pairwise Stitching plugin in Fiji (Schindelin et al., 2012). Higher-magnification images were acquired using 25×/0.8 NA water-immersion objective (LD LCI Plan-Apochromat 25×/0.8, 1 mm Corr, DIC M27) or 40×/1.1 NA water-immersion objective (LD C-Apochromat 40×/1.1 W Korr M27) objectives, and z-stacks encompassing the entire cell population of interest were collected. Images were acquired with an axial step size of 3.0 µm and a pixel dwell time of 0.728 µs. Low-magnification whole-larva images were acquired using the 20×/1.0 NA water-dipping objective with 0.7× digital zoom, whereas higher-magnification images were acquired using the 40×/1.1 NA water-immersion objective with 1.5× digital zoom, unless otherwise indicated.

For each experiment, imaging parameters were optimized using HCR-Cat, HCR-Immuno, or HCR-Multi samples to avoid pixel saturation and then applied to all samples. Separate detector gain settings were used for low- and high-magnification images. For experiments using Alexa Fluor 546, images were acquired only at 10%, 50%, and 100% laser power to minimize photobleaching. For all other experiments, detector gain was held constant while images were acquired at 0%, 0.5%, 1%, 2%, 5%, 10%, 25%, 50%, 75%, and 100% laser power.

For quantitative analysis, z-stacks acquired at different laser powers were concatenated in increasing order using the Concatenate plugin in Fiji, and the display range of the combined stack was reset to 65,535. Images were then converted to 8-bit, and the freehand selection tool was used to define regions of interest (ROIs) encompassing the labeled cell population. Integrated density and ROI area were measured for each z-plane. In samples where no cells were visible at the acquisition settings, the image display range was adjusted in Fiji solely to facilitate manual delineation of cell boundaries. The resulting ROIs were then applied to the original unmodified images for quantitative measurements. Background fluorescence was measured from a nearby region lacking labeled cells. For each laser power, integrated density values were summed across the entire z-stack for both the ROI and background regions, and the net fluorescence signal was calculated by subtracting the total integrated density of the background region from that of the ROI. Graphs were prepared using Prism (GraphPad). Absolute fluorescence intensity values were compared only among samples acquired within the same experiment under matched imaging conditions and were not used for quantitative comparisons across independently performed experiments.

### Puncta matching and correlation analysis

Images used for transcript-resolution analysis were acquired using a 40×/1.1 NA water-immersion objective (LD C-Apochromat 40×/1.1 W Korr M27) in Fast Airyscan mode with 5× digital zoom, yielding an XY pixel size of 0.0625 µm/pixel and an axial step size of 0.289 µm. Images were acquired with a pixel dwell time of 0.58 µs. Puncta were identified separately for the two *dbh* probe subsets in the imaging channels C1 (probe set 1, Alexa Fluor 546 for HCR v3.0, Alexa Fluor 568 for HCR-Immuno) and C2 (probe set 2, Alexa Fluor 633 for both HCR v3.0 and HCR-Immuno) using Fiji/ImageJ, and area, centroid coordinates, and integrated density were measured for each punctum. Object-based C1/C2 matching was performed using the geometric centroid coordinates (X/Y) for each punctum. Pairwise Euclidean distances between C1 and C2 puncta were calculated as √[(X_C2 − X_C1)² + (Y_C2 − Y_C1)²], and for each C1 punctum, the nearest C2 punctum was identified as the C2 object with the smallest centroid-to-centroid distance. Matches were classified as strong when the nearest C2 punctum was ≤5 pixels from the C1 punctum, and the C2 punctum with the minimum centroid-to-centroid distance was identified as the nearest-neighbor match. Matches were classified as possible when the nearest C2 punctum was >5 to ≤10 pixels from the nearest C1 punctum, and poor/no reliable match when the nearest C2 punctum was >10 pixels away. Only strong and possible matches were included in subsequent correlation analyses. For matched puncta, corresponding C1 and C2 puncta area and integrated density values were plotted as XY correlation graphs, with C1 values on the x-axis and C2 values on the y-axis. Correlations were assessed in Prism (GraphPad) using two-tailed Spearman correlation analysis. Spearman r, P values, and the number of matched puncta analyzed are reported for each correlation. Unimodality of puncta area and integrated fluorescence intensity distributions was assessed in R using Hartigan’s dip test for multimodality, with statistical significance defined as P < 0.05; distributions with P ≥ 0.05 were considered to show no significant departure from unimodality.

### Adult zebrafish brain sample preparation

Adult zebrafish brain HCR was performed according to a modified version of a published protocol (Kenney et al., 2021). Adult zebrafish (2–3 years old) were anesthetized in 4% tricaine and euthanized by immersion in ice-cold water for 5 min. Animals were decapitated, and the heads were incubated in ice-cold 1× DPBS for 5 min to allow blood to drain. The dorsal skin and skull were removed to expose the brain, after which heads were fixed overnight at 4°C in 4% PFA in DPBST. Brains were subsequently dissected in ice-cold 1× DPBS and fixed overnight a second time in ice-cold 4% PFA in DPBST.

Unless otherwise indicated, all incubations and washes were performed with gentle shaking, whereas dehydration and rehydration steps were performed without shaking. Probe hybridization and wash steps at 37°C were carried out using a heat block. A short segment of the spinal cord was retained to facilitate sample handling and mounting for imaging. Brains were washed three times for 5 min in DPBSTx, dehydrated through a graded methanol series (20%, 40%, 60%, 80%, and 100% methanol; 30-60 min per step) at room temperature, incubated in ice-cold 100% methanol on ice for 1 h, and then transferred to chilled 5% hydrogen peroxide in methanol overnight at 4°C. Samples were rehydrated through a reverse methanol series (80%, 60%, 40%, and 20% methanol; 30-60 min per step) at room temperature, washed once in 1× DPBS, and washed twice for 1 h in DPBSTx. Permeabilization was performed overnight at 37°C in permeabilization solution (0.3 M glycine and 20% DMSO in DPBSTx), followed by three 5-min washes in DPBSTx. Typically, 2–5 brains were processed in 2.0 mL microcentrifuge tubes, using 500–750 μL of solution for each incubation and wash step.

### HCR v3.0 for adult zebrafish brain

After adult brain sample preparation, brains were processed using the HCR workflow described for larval zebrafish, except that the probe concentration was increased to 20 nM and probe hybridization was extended to 36-60 h. HCR v3.0 samples used as controls for HCR-Cat experiments were stored in 5× SSCT at 4°C in the dark until the HCR-Cat-specific steps were completed.

After labeling, brains were dehydrated through a graded methanol series (20%, 40%, 60%, 80%, and 2× 100% methanol; 30-60 min per step), followed by overnight incubation in 100% methanol at room temperature. Samples were then incubated in 66% dichloromethane (DCM; Millipore Sigma, 270997) in methanol for 3 h at room temperature, washed twice for 15 min in 100% DCM without shaking, and transferred to dibenzyl ether (DBE; Millipore Sigma, 33630) for storage at 4°C in the dark until imaging.

### HCR-Cat for adult zebrafish brain

After tissue preparation, brains were processed using the HCR workflow and HCR-Cat protocol for larval zebrafish, except that samples were blocked for 2 h in Roche Blocking Solution and incubated with anti-Flu-HRP (Roche, 11426346910; 1:400) for 36-60 h at 4°C. Samples were washed six times for 15 min in DPBSTx, and signal was developed for 45 min at room temperature using TSA PLUS Fluorescein diluted 1:200 in TSA amplification diluent. Following signal development, samples were washed five times for 5 min in DPBSTx and incubated overnight in DPBSTx at 4°C to reduce background fluorescence. Brains were subsequently dehydrated and cleared as described above for HCR v3.0 samples.

### Imaging of adult zebrafish brain

Cleared brains were mounted using Norland Optical Adhesive 61 (Norland Optical). A cured base layer of adhesive was first prepared on a sample holder compatible with the light-sheet imaging chamber, followed by a second drop into which the brain was positioned. The retained spinal cord was used to orient the brain before curing the adhesive, thereby securing the sample in the desired orientation. Brains were imaged in DBE using a LaVision BioTec Ultramicroscope II light-sheet microscope at 2× magnification.

### Statistical analysis

Statistical analyses were performed using Prism (GraphPad). Data are presented as mean ± SEM unless otherwise indicated. Comparisons between multiple experimental groups were performed using one-way or two-way ANOVA, as appropriate, followed by multiple-comparisons testing. Correlations between matched puncta were assessed using two-tailed Spearman correlation analysis. The statistical tests used for each experiment are specified in the corresponding figure legend. P < 0.05 was considered statistically significant. Information on statistical tests and results are provided in Table S1.

### Mice

#### Mouse husbandry

Mouse husbandry and all experimental procedures were performed in accordance with the Guide for the Care and Use of Laboratory Animals of the National Institutes of Health and approved by the Institutional Animal Care and Use Committee (IACUC; protocol 1746) and the Office of Laboratory Animal Resources at the California Institute of Technology. Eight-week-old male C57BL/6J (strain #000664) and Thy1-YFP-H (strain #003782) mice were obtained from The Jackson Laboratory. Mice were housed in groups of 3–4 per cage under a 12 h light/12 h dark cycle with ad libitum access to food and water.

#### Mouse tissue processing

Mice were euthanized by intraperitoneal injection of Euthasol (100 mg/kg) and transcardially perfused with 30 mL of ice-cold heparinized 1× PBS, followed by 30 mL of ice-cold 4% paraformaldehyde (PFA) in 1× DPBS. Whole intact skulls were post-fixed in 4% PFA in 1× DPBS at 4°C for at least 48 h. Skulls were washed three times in 1× PBS, after which brains were dissected and stored in 1× DPBS at 4°C until further processing. For 100 μm sections, brains were cryoprotected by immersion in 30% sucrose in 1× DPBS until the tissue had sunk, then embedded in O.C.T. Compound (Scigen, #4586), flash-frozen in a dry ice–ethanol bath, and stored at −70°C until sectioning. Coronal sections were cut using a cryostat (Leica Biosystems), collected in 1× DPBS, and stored at 4°C until use. For 600 μm sections, brains were sectioned coronally using a vibrating microtome (Leica Biosystems), collected in ice-cold 1× DPBS, and stored at 4°C until use.

#### PACT tissue clearing

Thick (600 μm) coronal mouse brain sections were PACT-cleared according to published protocols (Treweek et al., 2015; Yang et al., 2014). Briefly, sections were incubated overnight at 4°C in a hydrogel solution consisting of 1× DPBS, 4% acrylamide (Bio-Rad, #1610140), 1% PFA, and 0.25% VA-044 thermal polymerization initiator (Wako, VA-044). Oxygen was removed by bubbling nitrogen gas through the solution for 5 min, after which the hydrogel was polymerized for 2 h at 37°C. Excess hydrogel was removed by briefly washing the tissue in 1× DPBS, followed by delipidation in 1× DPBS containing 8% SDS for 16 h. Samples were then washed in 1× DPBS with six 1-h washes using excess buffer (50 mL per wash).

#### Common HCR workflow for mouse tissue

Split-initiator probe sets were designed using the ProbeDesignSplit workflow (https://github.com/GradinaruLab/HCRprobe) according to published methods (Jang et al., 2023), based on the HCR v3.0 amplifier sequences described by (Choi et al., 2018). Candidate probes were selected based on GC content, sequence complexity, predicted off-target hybridization, thermodynamic stability, and cross-dimerization as implemented in the ProbeDesignSplit workflow. Probe sets were ordered from Integrated DNA Technologies and diluted in Ultrapure water before use. Sequences of the mouse split-probe sets designed for this study are provided in Table S2. Unless otherwise indicated, all incubations were performed at room temperature.

For 100 μm sections, tissue was mounted on glass slides (Brain Research Laboratories, #2575-plus) and permeabilized for 1 h in 1× DPBS containing 0.1% Triton X-100 (DPBSTx). PACT-cleared 600 μm sections were stained free-floating. Unless otherwise indicated, 100 μm and PACT-cleared sections were processed identically.

Samples were equilibrated for 1 h at 37°C in hybridization buffer consisting of 2× SSC, 10% ethylene carbonate, and 10% dextran sulfate, followed by hybridization for 12-16 h at 37°C in pre-warmed hybridization buffer containing 2 nM of each probe. Slide-mounted sections were covered with a coverslip and incubated in a humidified chamber to prevent drying. Following hybridization, samples were washed twice for 30 min in stringent wash buffer (2× SSC, 30% ethylene carbonate) at 37°C, followed by two 30-min washes in 5× SSCT at room temperature. Slide-mounted sections were washed using approximately 1 mL per wash of stringent wash buffer pipetted directly onto the slide, then 60 mL per wash of 5× SSCT in a Coplin jar. PACT-cleared sections were washed in 5 mL of each buffer (up to three sections per tube).

Samples were then equilibrated in HCR amplification buffer (2× SSC, 10% ethylene carbonate) for 1 h. Alexa Fluor 647-, FITC-, or DIG-conjugated HCR amplifier hairpins were snap-cooled individually by heating to 95°C for 90 s followed by passive cooling at room temperature in the dark for 30 min. Amplifier hairpins were then combined, diluted to a final concentration of 60 nM in amplification buffer, and incubated with the tissue overnight in the dark. Slide-mounted sections were again covered with a coverslip and maintained in a humidified chamber.

Following HCR amplification, samples were washed twice for 30 min in 5× SSCT, followed by three 15-min washes in DPBSTx. At this point, the protocol diverged according to the signal detection strategy described below.

#### HCR v3.0 for mouse tissue

After the common HCR workflow for mouse tissue, HCR v3.0 samples were stored in 5× SSCT at 4°C until HCR-Cat, HCR-Immuno, and HCR-Multi samples had completed processing. Samples were then processed for nuclear staining and mounting as described below.

#### HCR-Cat for mouse tissue

After the common HCR workflow for mouse tissue, tissue labeled with FITC-conjugated HCR amplifier hairpins was blocked for 1 h in TNB blocking solution (Akoya Biosciences, FP1012), followed by overnight incubation at 4°C with sheep anti-Flu-HRP Fab fragments (Roche, 11426346910; 1:100) diluted in TNB blocking solution. Samples were washed three times for 30 min in DPBSTx. Signal in 100 μm sections was developed for 10 min using the Alexa Fluor 488 Tyramide SuperBoost Kit (Thermo Fisher, B40932) according to the manufacturer’s instructions. Signal in PACT-cleared sections was developed for 10 min using an Alexa Fluor 647 Tyramide SuperBoost Kit (Thermo Fisher, B40936) according to the manufacturer’s instructions. Following signal development, the reaction was terminated, and samples were washed four times for 30 min in DPBSTx, followed by an overnight incubation in DPBSTx. Samples were then processed for nuclear staining and mounting as described below.

#### HCR-Immuno for mouse tissue

After the common HCR workflow, tissue labeled with FITC- or DIG-conjugated HCR amplifier hairpins was blocked for 1 h in DPBSTx containing 10% normal donkey serum (Jackson ImmunoResearch, 017-000-121). Samples were incubated overnight at 4°C with either sheep anti-Flu (SouthernBiotech, 6400-01; 1:100) or rabbit anti-DIG (Invitrogen, 700772; 1:100) diluted in the same blocking buffer. Samples were washed five times for 30 min in DPBSTx, followed by overnight incubation at 4°C with Alexa Fluor 647-conjugated donkey anti-sheep F(ab’)_₂_ or donkey anti-rabbit F(ab’)_₂_ secondary antibodies (Jackson ImmunoResearch, 713-606-147 and 711-606-152, respectively; 1:200) diluted in DPBSTx containing 10% normal donkey serum. Samples were washed five additional times for 30 min in DPBSTx before nuclear staining and mounting as described below.

#### HCR-Multi for mouse tissue

After the common HCR workflow for mouse tissue, tissue labeled with DIG-conjugated HCR amplifier hairpins was blocked for 1 h in antibody buffer (Molecular Technologies), followed by overnight incubation with rabbit anti-DIG (Invitrogen, 700772; 1:100) diluted in antibody buffer. Samples were washed five times for 30 min in DPBSTx, followed by overnight incubation with B2 initiator-conjugated donkey anti-rabbit secondary antibody (Molecular Technologies; 1:500). Samples were washed five additional times for 30-min in DPBSTx, equilibrated in HCR amplification buffer for 1 h, and subjected to a second overnight HCR amplification using FITC-conjugated B2 hairpins (60 nM) prepared as described above. Following amplification, samples were washed twice for 30 min in 5× SSCT, followed by three 30-min washes in DPBSTx, and were subsequently processed as described above for HCR-Immuno using FITC-conjugated amplifier hairpins before nuclear staining and mounting.

#### Final processing

Following completion of the staining procedures, mouse samples were stained with Hoechst 33342 (Thermo Fisher, H3570; 1:10,000) in 1× DPBS for 20 min, followed by three 10-min washes in 1× DPBS. PACT-cleared sections underwent an additional wash in 0.02 M phosphate buffer. For 100 μm sections, coverslips were mounted using ProLong Diamond (Thermo Fisher, P36970). PACT-cleared sections were incubated in 5 mL of refractive index matching solution (RIMS) consisting of 40 g iohexol (Histodenz; Sigma-Aldrich, D2158) dissolved in 30 mL of 0.02 M phosphate buffer (Electron Microscopy Sciences, 19340-72; pH 7.4; refractive index 1.46).

Once the sections became optically transparent, they were mounted in RIMS using iSpacer mounting chambers (SunJin Lab, IS011) and coverslips.

#### Imaging of mouse samples

Stained mouse sections were imaged using a Zeiss LSM 880 confocal microscope with a Plan-Apochromat 10×/0.45 NA air objective. Images for *Snap25*, *Eno2*, *Slc1a3*, and *Th* are single planes, acquired with a pixel dwell time of 0.41 μs and an axial resolution of 4.98 μm (step size: 2.49 μm). Images of *Drd1*, *Drd2*, *Tacr1*, and *Tacr3* are maximum intensity projections of 7 planes acquired with a pixel dwell time of 1.02 μs and an axial resolution of 4.98 μm (step size: 2.49 μm). PACT-cleared mouse brain slices were imaged in 178 planes acquired with a pixel dwell time of 0.77 μs and an axial resolution of 6.82 μm (step size: 3.41 μm). Imaging settings were optimized to capture the full dynamic range of the fluorescence signal without pixel saturation. Whenever possible, laser power was adjusted before detector gain. 633 nm laser powers, expressed as percent of maximal output, used for imaging RNA signal in PACT-cleared tissue sections are reported on figures. 488 nm laser powers used for imaging RNA signal in 100 μm sections are: 5% (*Snap25*), 10% (*Eno2*), 15% (*Drd1*, *Drd2*, *Tacr1*, and *Tacr3*), 20% (*Slc1a3*), and 100% (*Th*). Fields of view were selected using non-experimental channels (e.g. Hoechst or YFP) to avoid selection bias

#### Image analysis of mouse samples

For analysis of PACT-cleared mouse tissue, 600 μm confocal stacks were subdivided into thirty 23 μm blocks, and every second block was analyzed. For HCR-Cat, YFP-positive cell bodies were segmented using the Trainable Weka Segmentation plugin in Fiji, and the resulting regions of interest (ROIs) were used to quantify RNA fluorescence intensity. For HCR-Immuno and HCR-Multi, YFP-positive cell bodies were segmented using Cellpose-SAM (version 4.0.8; (Pachitariu et al., 2025)), and RNA fluorescence intensity was quantified using CellProfiler (version 4.2.8.1; (Stirling et al., 2021)).

### Statistical analysis

Statistical analyses were performed using Prism (GraphPad). The statistical tests used for each experiment are specified in the corresponding figure legends. P < 0.05 was considered statistically significant. Information on statistical tests and results are provided in Table S1.

#### Drosophila

##### Drosophila husbandry

*Drosophila* experiments were performed in accordance with California Institute of Technology Institutional Animal Care and Use Committee guidelines. All experiments were performed using wandering third-instar *Drosophila melanogaster* larvae (Canton S), which were maintained at room temperature under standard laboratory conditions.

##### HCR v3.0 staining of *Drosophila* samples

HCR reagents, including probes and HCR amplifier hairpins (B1-B5) (Choi et al., 2018), were purchased from Molecular Technologies and Molecular Instruments. The *Drosophila* probe sets used in this study were commercially designed and are available for purchase from Molecular Technologies. Probe sets may also be designed using published scripts (Ćorić et al., 2023; Olivares-Abril et al., 2024). Staining was performed according to a modified version of a published protocol (Duckhorn et al., 2022) using ethylene carbonate-based hybridization buffers (all % v/v): Hybridization buffer: 2× SSC, 10% ethylene carbonate, 10% dextran sulfate, and 0.1% Tween-20; Stringent wash buffer: 2× SSC, 30% ethylene carbonate, and 0.1% Tween-20; Wash buffer: 2× SSC and 0.1% Tween-20; Amplification buffer: 2× SSC, 10% ethylene carbonate, and 0.1% Tween-20.

Dissections were performed as previously described (Menon et al., 2015). Briefly, wandering third-instar larvae were dissected in 1× DPBS, and brain lobes together with ventral nerve cords (VNCs) were fixed in 4% PFA in DPBST for 30 min at room temperature, followed by three 5-min washes in DPBSTx. Typically, 10-12 samples were processed in 1.5-mL microcentrifuge tubes using 500 μL of solution per incubation and wash step. Unless otherwise indicated, all incubations and washes were performed with gentle shaking on a nutator, whereas incubations at 37°C were performed using a heat block.

After tissue preparation, samples were processed using the common HCR workflow for larval zebrafish, except that the ethylene carbonate-based hybridization buffer system described above was used throughout. Probe hybridization was performed using 16 nM probes for weakly expressed transcripts or 2-4 nM probes for highly expressed transcripts. For weakly expressed targets, probe hybridization was extended to 36-60 h. HCR v3.0 control samples were stored in 2× SSCT at 4°C in the dark until the HCR-Cat-specific steps were completed. Following labeling, samples were washed once in 1× DPBS for 10 min without shaking, mounted in 100% VECTASHIELD PLUS (50-80 μL) until imaged.

##### HCR-Cat staining of *Drosophila* samples

After tissue preparation, samples were processed according to the common HCR workflow for larval zebrafish and the HCR-Cat protocol for larval zebrafish, except that the ethylene carbonate-based hybridization buffer system described above was used throughout. Following removal of excess HCR amplifier hairpins, samples were washed three times for 5 min in DPBSTx, blocked for at least 1 h in Roche Blocking Solution, and incubated overnight at 4°C with anti-Flu-HRP Fab fragments (Roche, 11426346910; 1:400). Samples were washed five times for 15 min in DPBSTx before signal development for 30 min at room temperature using TSA PLUS Fluorescein diluted 1:200 for weakly expressed targets or 1:300 for highly expressed targets. Following signal development, samples were washed five times for 5 min in DPBSTx, followed by one 10-min wash in 1× DPBS, mounted in 100% VECTASHIELD PLUS, and imaged as described below. For all experiments, samples were incubated overnight in DPBSTx at 4°C before mounting to reduce background fluorescence.

##### Imaging of *Drosophila* samples

*Drosophila* samples were mounted in 100% VECTASHIELD PLUS and imaged using a Zeiss LSM 880 confocal microscope with either a 20×/1.0 NA water-dipping objective (W Plan-Apochromat 20×/1.0 DIC CG = 0.17 M27, 75 mm) or a 40×/1.1 NA water-immersion objective (LD C-Apochromat 40×/1.1 W Korr M27) objective. Z-stacks were acquired with axial step sizes of 0.659 µm and 0.923 µm for the 20× and 40× imaging configurations, respectively. Images were acquired with a pixel dwell time of 0.728 µs.

## Acknowledgements

We thank Alex Mack, Caressa Wong, Barbara Orozco, and Axel Dominguez for zebrafish care; Drs. Andres Collazo and Zhongying Wang at the Biological Imaging Center at Caltech for help with imaging; and all members of the Prober, Gradinaru, and Zinn labs for comments and suggestions.

## Data and Resource Availability

All relevant data and details of resources can be found within the article and its supplementary information.

## Author Contributions

Conceptualization: CS, DAP Methodology: CS, NB, GC, JX, GS, YS

Investigation: CS, NB, GC, JX, JYP, UH, MSG, TC Visualization: CS, NB, GC, JX, DAP

Funding acquisition: VG, KZ, DAP Project administration: CS, VG, KZ, DAP Supervision: CS, VG, KZ, DAP

Writing – original draft: CS, DAP

Writing – edit and revision: CS, NB, GC, JX, UH, VG, KZ, DAP

