## Supplementary Figures 1-9 for "Hybridization chain reaction variants with enhanced sensitivity for detecting challenging mRNA targets"

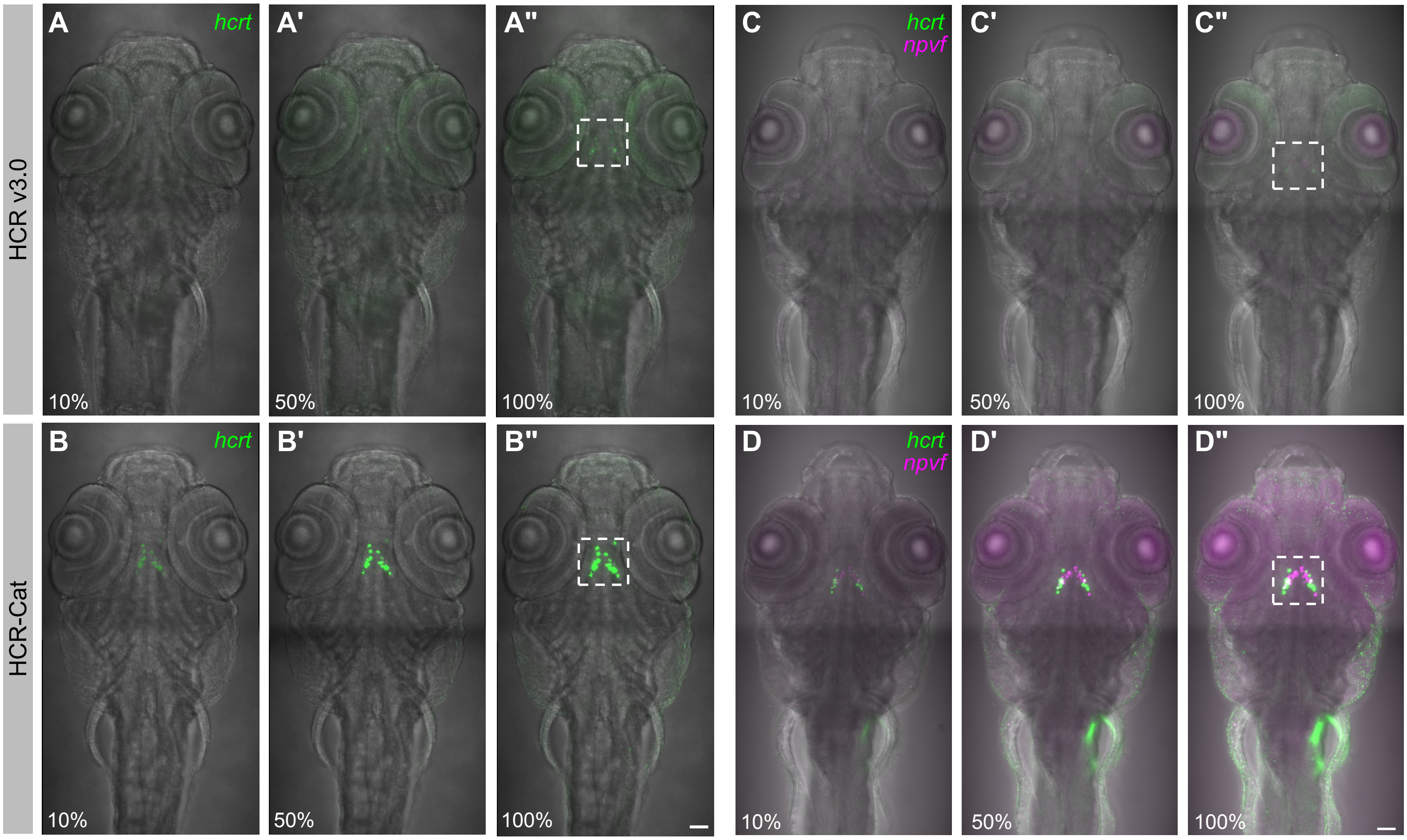


**Figure S1: HCR-Cat with FITC enhances mRNA detection sensitivity of *hcrt* and *npvf* compared to HCR v3.0 in zebrafish larvae. (A–B)** Detection of *hcrt* with 8 probes showed significantly higher signal using HCR-Cat compared to HCR v3.0. **(C–D)** Co-detection of *hcrt* and *npvf* with 8 and 10 probes, respectively, showed significantly higher signals for both targets using HCR-Cat with FITC-conjugated amplifiers for *hcrt* and DIG-conjugated amplifiers for *npvf* compared to HCR v3.0. The same region was imaged at 10%, 50% and 100% laser power while maintaining a fixed detector gain. The boxed regions are shown at a higher magnification in **Fig. 1.** A higher digital gain was used to detect *hcrt* in (**A–B**) compared to (**C-D**). All experiments used at least 4 fish. Representative images shown are maximum intensity projections of z-stacks across the entire cell populations. Alexa Fluor 488 **(A, C)**, Alexa Fluor 546 **(C)**, TSA-Flu **(B, D)**, and TSA-Cy3 **(D)** were used. Scale bars: 50 µm.

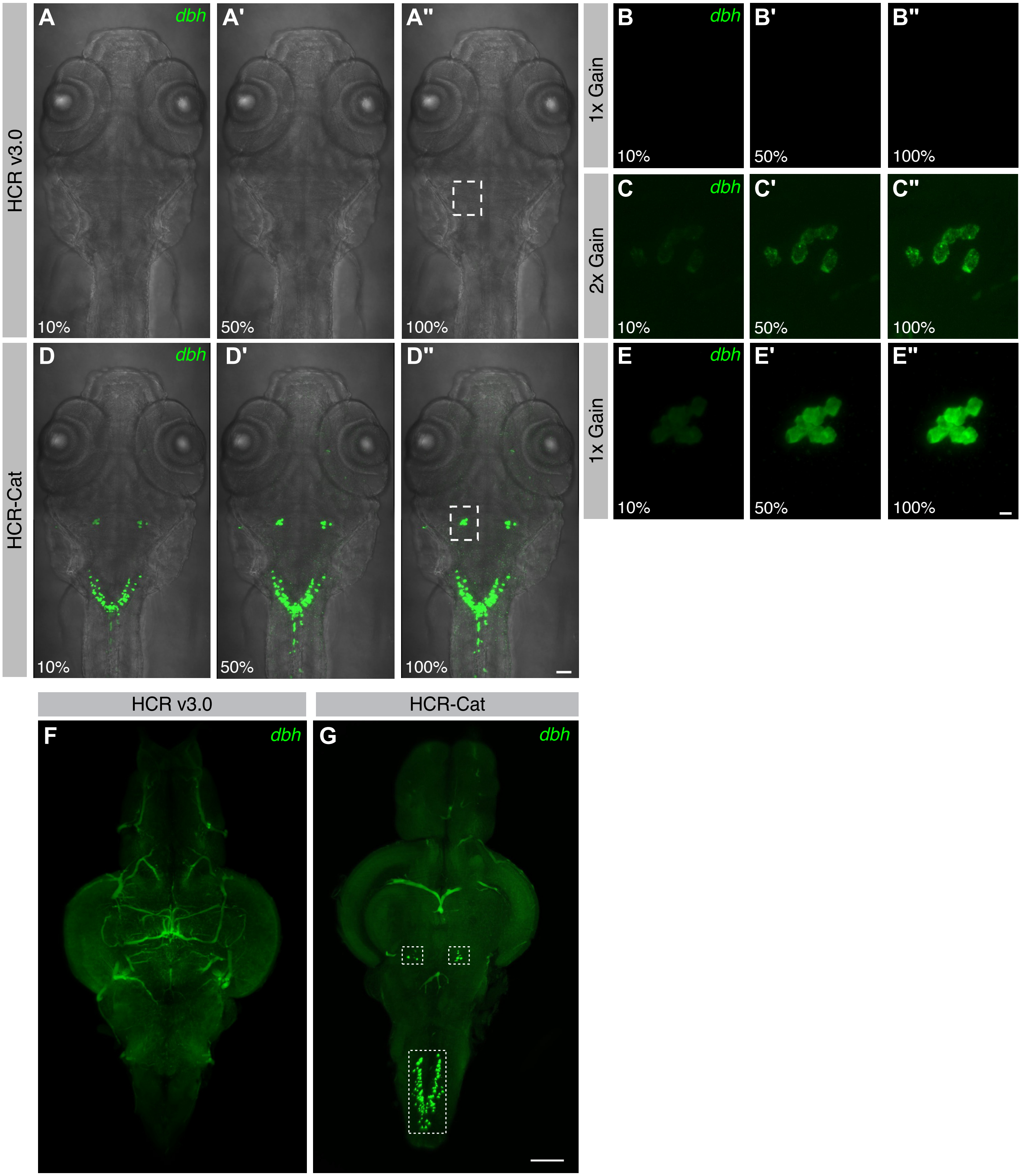


**Figure S2: HCR-Cat with FITC enhances *dbh* mRNA detection sensitivity compared to HCR v3.0 in larval and adult zebrafish brain. (A–E")** Detection of *dbh* using 20 probes with HCR v3.0 **(A–C")** or HCR-Cat with FITC-conjugated amplifiers **(D–E")**. The same region was imaged at 10%, 50% and 100% laser power for all samples while maintaining a fixed detector gain, except for **(C–C")**, where a two-fold higher digital gain was used. The boxed regions in **(A")** and **(D")** are shown at higher magnification in **(B–B")**, **(C–C")**, and **(E–E")**. **(F-G)** Detection of *dbh* using 20 probes in a whole-mount adult zebrafish brain with HCR v3.0 **(F)** and HCR-Cat **(G)**. Using the same laser power and detector gain, HCR-Cat produced a robust signal, whereas HCR v3.0 failed to yield detectable transcript signal. The boxed regions indicate *dbh* expression in the locus coeruleus (squares) and medulla oblongata (rectangle). The other green structures are auto-fluorescent blood vessels. Each experiment used at least 4 fish. Representative images shown are maximum intensity projections of z-stacks across the entire *dbh* cell populations. Alexa Fluor 488 (**A–C", F**), and TSA-Flu (**D–E", G**) were used. Scale bars: 50 µm (**D"**), 10 µm (**E"**) and 100 µm (**G**).

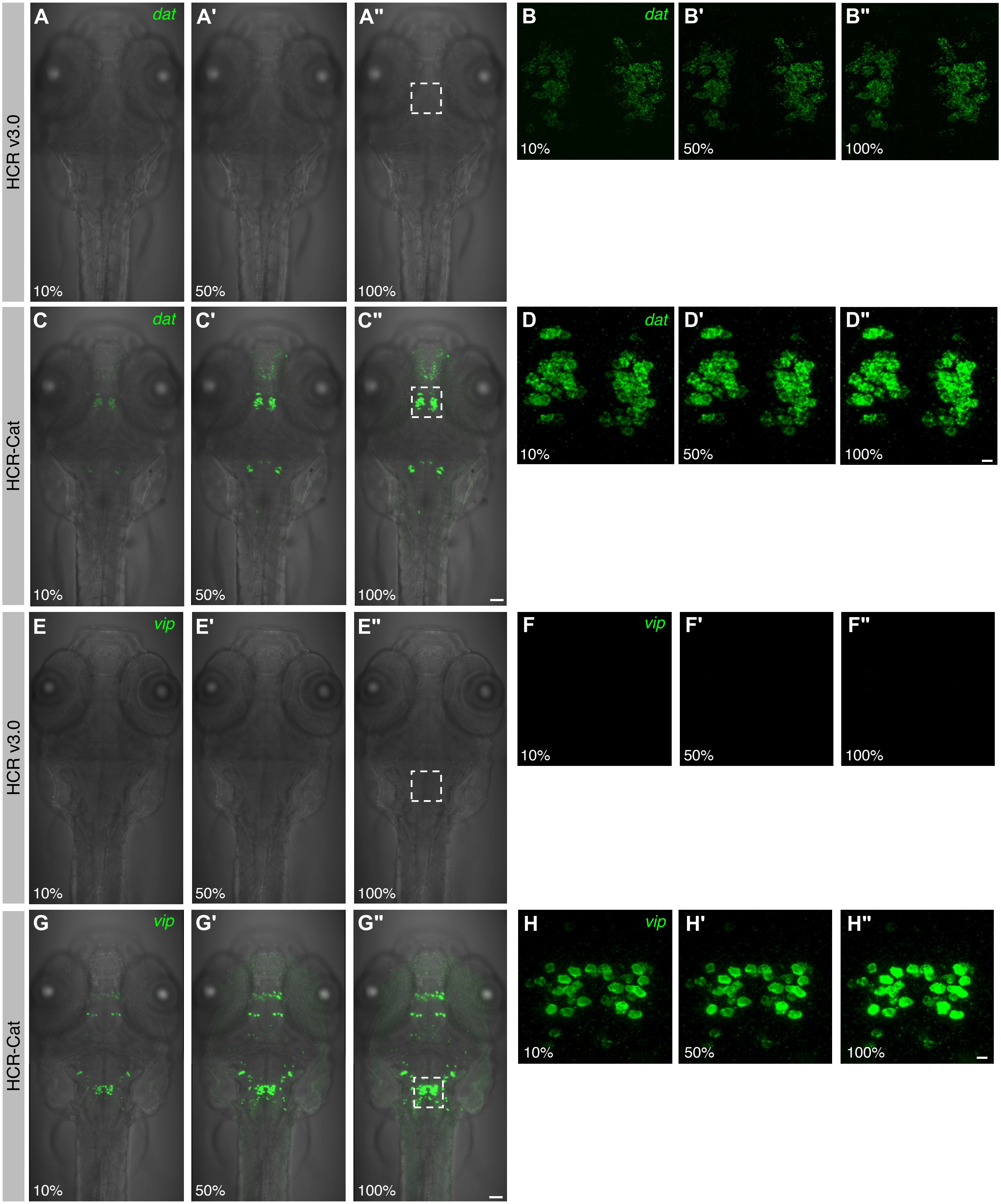


**Figure S3: HCR-Cat with FITC enhances *dat* and *vip* mRNA detection sensitivity compared to HCR v3.0 in zebrafish larvae. (A–D")** Detection of *dopamine transporter* (*dat*) using 20 probes with HCR v3.0 (**A–B"**) or HCR-Cat with FITC-conjugated amplifiers (**C–D"**). **(E–H")** Detection of *vasoactive intestinal peptide* (*vip*) using 11 probes with HCR v3.0 (**E–F"**) or HCR-Cat with FITC-conjugated amplifiers (**G–H"**). The same regions were imaged at 10%, 50%, and 100% laser power while maintaining a fixed detector gain. The boxed regions in (**A"**), (**C"**), (**E"**) and (**G"**) are shown at higher magnification in (**B–B"**), (**D–D"**), (**F–F"**) and (**H–H"**). All experiments used at least 4 fish. Representative images shown are maximum intensity projections of z-stacks across the entire *dat* and *vip* cell populations. Alexa Fluor 488 (**A–B", E–F"**) and TSA-Flu (**C–D", G–H"**) were used. Scale bars: 50 μm (**C", G"**) and 10 μm (**D", H"**).

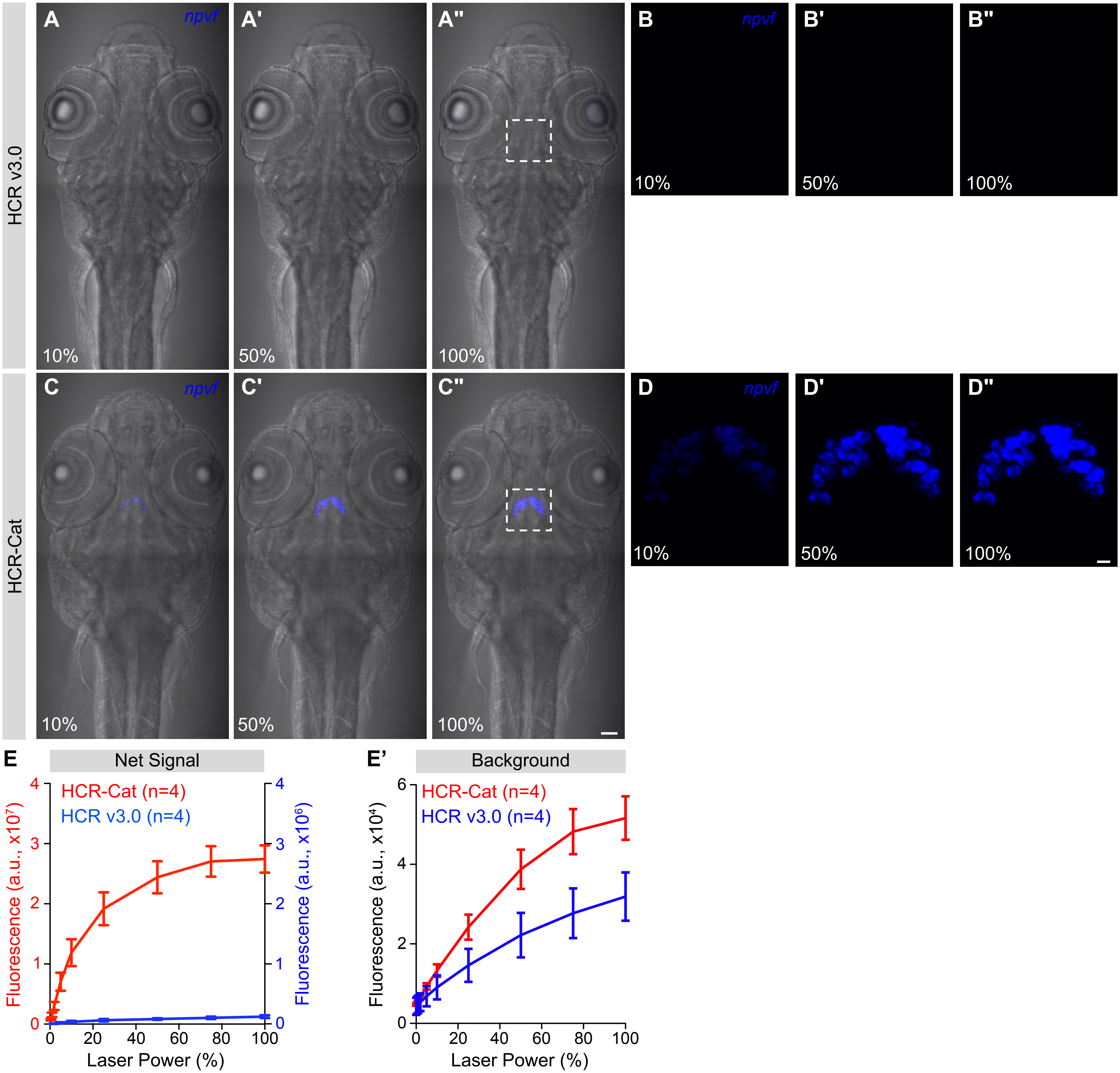


**Figure S4: HCR-Cat with DIG enhances *npvf* mRNA detection sensitivity compared to HCR v3.0 in zebrafish larvae. (A–D)** Detection of *npvf* using 10 probes with HCR v3.0 (**A–B"**) or HCR-Cat with DIG-conjugated amplifiers (**C–D"**). Alexa Fluor 647 (**A–B"**) and TSA-Cy5 (**C–D"**) were used. The boxed regions in (**A"**) and (**C"**) are shown at higher magnification in (**B-B"**) and (**D-D"**). For each sample, the boxed region in (**A",C"**) was imaged by gradually increasing the laser power from 0% to 100% while maintaining a fixed detector gain. Scale bars: 50 µm (**C"**) and 10 µm (**D"**). **(E)** Quantification of net signal using *npvf* probes and either HCR v3.0 (blue, n=4 fish) and HCR-Cat (red, n=4 fish). HCR-Cat produced significantly greater net signal than HCR v3.0 at laser powers of 5% and above ($p < 0.001$ at 5% laser power, $p < 0.0001$ at all higher laser powers). There was no significant difference in HCR-Cat net signal between 50% and higher laser powers, indicating that the net signal approached saturation at 50% laser power. (**E′**) Quantification of background fluorescence used for calculation of net signal in (**E**). HCR-Cat showed significantly higher background than HCR v3.0 at laser powers of 50% and above ($p < 0.01$ at 50% laser power, $p < 0.001$ at 75% and 100% laser powers), with no significant differences at lower laser powers. All experiments used at least 4 fish. Mean ± SEM (arbitrary units, a.u.) are shown (**E, E'**). Representative images shown are maximum intensity projections of z-stacks across the entire cell populations. Scale bars: 50 µm (**C**) and 10 µm (**D"**). Statistical significance was assessed using Two-way ANOVA with amplification method and laser power as factors, followed by Tukey's multiple-comparisons test.

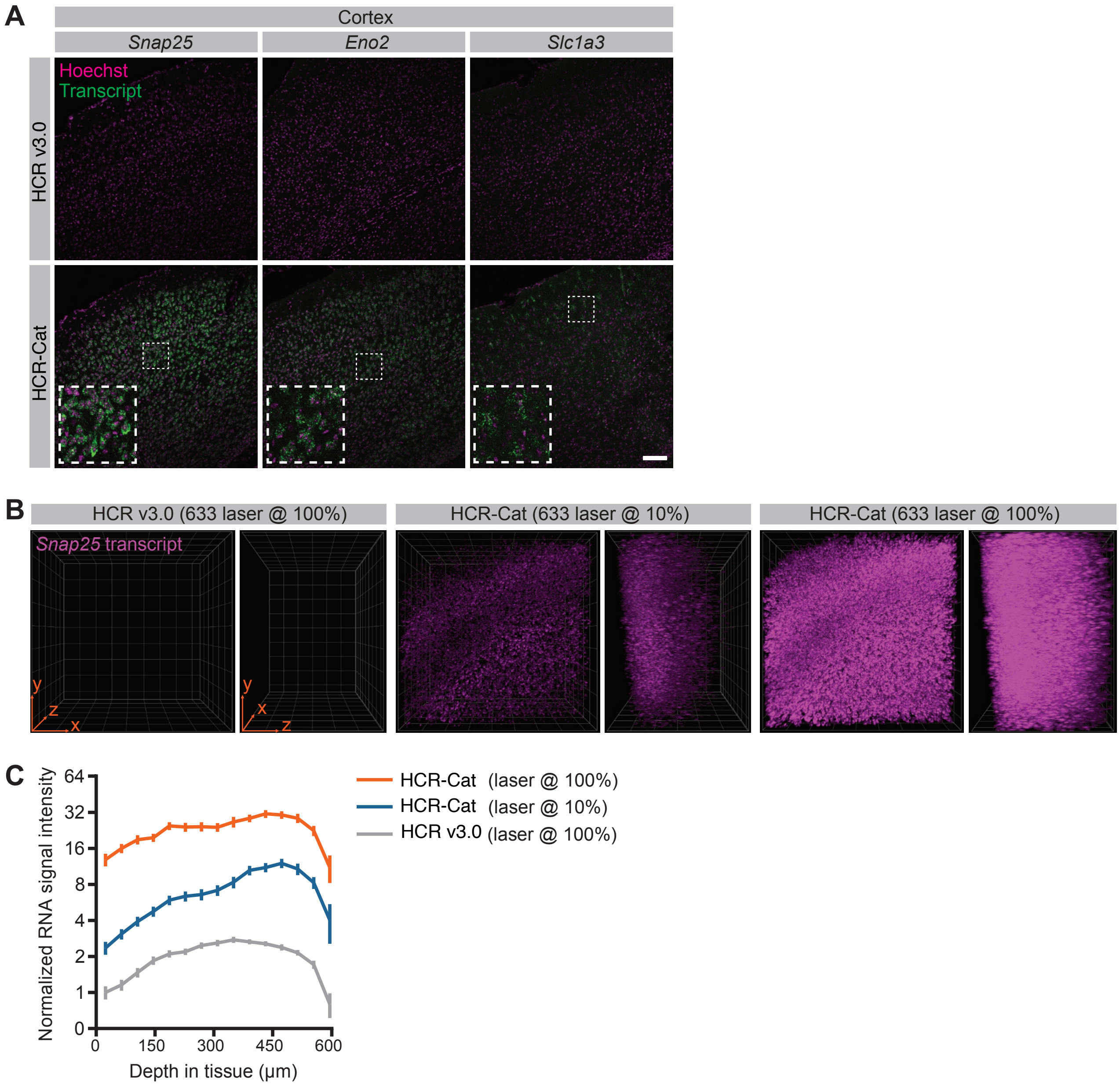


**Figure S5: HCR-Cat with FITC enhances mRNA detection sensitivity in mouse cortex. (A)** Comparison of HCR-Cat with HCR v3.0 for detection of the pan-neuronal markers *Snap25* (15 probes) and *Eno2* (19 probes), and the glial marker *Slc1a3* (12 probes), in mouse cortex. Each staining was performed on 100 µm-thick sections from 3 mice. Representative images shown are maximum intensity projections of z-stacks. Boxed regions are shown at higher magnification in each inset. **(B)** Confocal stacks through 600 µm of PACT-cleared Thy1-YFP mouse cortex showing detection of *Snap25* transcript with HCR v3.0 and HCR-Cat. HCR v3.0 samples were imaged at 100% laser power, whereas HCR-Cat samples were imaged at 10% or 100% laser power. No correction for imaging depth was applied. Representative images from 4 mice are shown. **(C)** Quantification of normalized *Snap25* mRNA signal intensity as a function of tissue depth. RNA signal was quantified in YFP-positive cell bodies segmented using the YFP channel (not shown) and normalized to the HCR v3.0 mRNA signal intensity at the tissue surface. HCR-Cat yielded an approximately 10-fold increase in *Snap25* mRNA signal compared to HCR v3.0 when imaged at the same laser power ($p < 0.001$, at all tissue depths). Decreasing the laser power to 10% still yielded an approximately 3-fold increase in *Snap25* mRNA signal compared to HCR v3.0 imaged at 100% laser power ($p < 0.05$ from 309.4 µm to 595 µm). Mean ± 95% confidence interval from 19-109 cells per depth, pooled from 4 mice, is shown. Statistical significance was tested with a mixed effects analysis with Dunnett's multiple comparison test, treating the animal as the unit of replication. Complete statistical results, including adjusted p-values for pairwise comparisons, are provided in Table S1. Alexa Fluor 488 (**A**, top panel), Alexa Fluor 647 (**B**, left panels), TSA-Alexa Fluor 488 (**A**, bottom panel), and TSA-Alexa Fluor 647 (**B**, middle and right panels) were used. Scale bars: 100 µm (**A**) and 160 µm (**B**, orange arrows in plane of image).

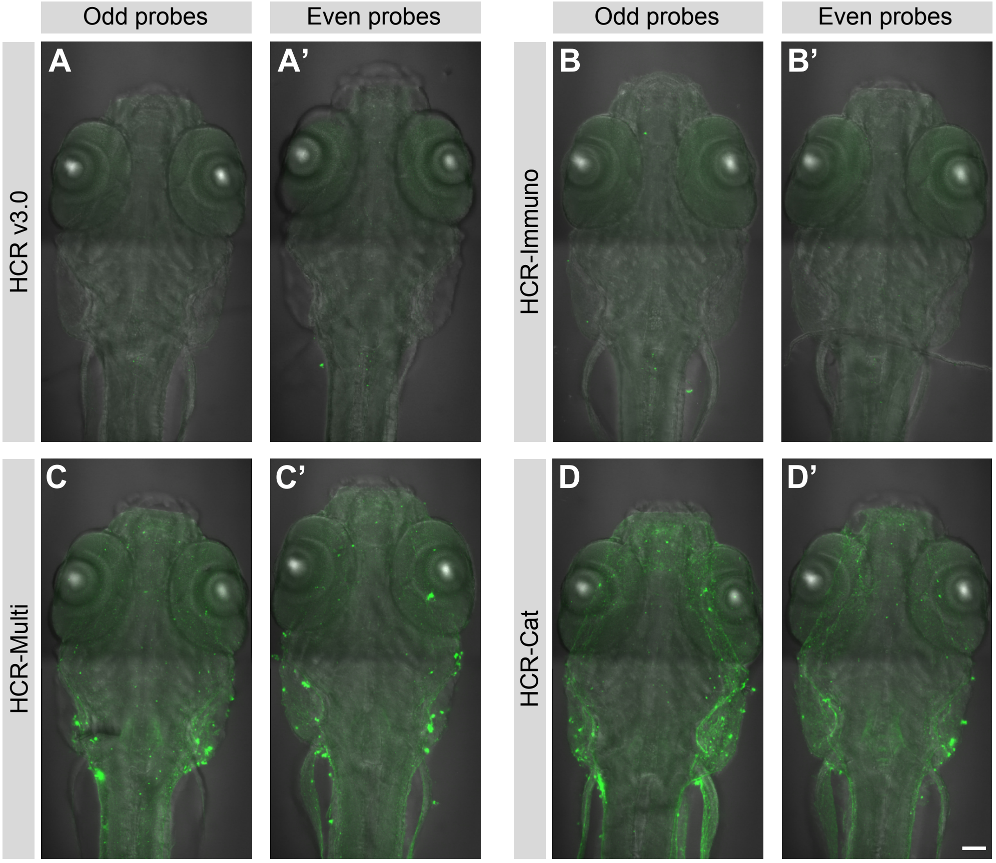


**Figure S6: Odd- and even-half-probe set controls confirm the specificity of HCR-Cat, HCR-Multi, and HCR-Immuno in zebrafish larvae.** HCR v3.0 (**A–A′**), HCR-Immuno (**B–B′**), HCR-Multi (**C–C′**), and HCR-Cat (**D–D′**) were performed using either the odd-half-probe set (**A–D**) or the even-half-probe set (**A′–D′**) for detection of *hcrt* mRNA in zebrafish larvae. None of the four methods produced specific labeling of *hcrt*-positive neurons when either half-probe set was used alone, confirming that signal generation requires adjacent binding of both members of each split-probe pair to generate a complete initiator. Occasional bright puncta were observed but these were mostly located outside the brain and did not exhibit the labeling pattern of *hcrt* neurons. Representative images shown are maximum intensity projections of z-stacks through the brain region where *hcrt* neurons are located. Alexa Fluor 488 (**A–C'**) and TSA-Flu (**D–D'**) were used. Scale bar: 100 μm.

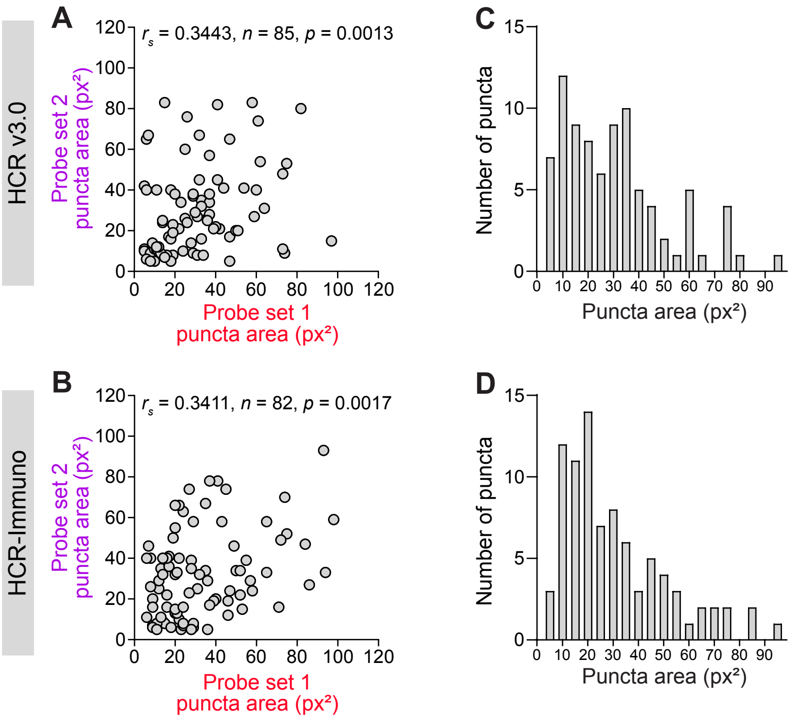


**Figure S7: Object-based analysis of HCR-Immuno *dbh* neuron matched puncta areas. (A–B)** Scatter plots show matched puncta areas for *dbh* probe subsets 1 (red) and 2 (purple) detected using HCR v3.0 (**A**) and HCR-Immuno (**B**). Matched puncta showed a positive and statistically significant correlation by Spearman correlation analysis (*r*□ ) for both HCR v3.0 and HCR-Immuno. n = number of puncta analyzed**. (C–D)** Frequency distribution of puncta area for HCR v3.0 (**C**) and HCR-Immuno (**D**). Both methods showed a predominant single population of similarly sized puncta with comparable puncta size distributions, indicating that the punctate morphology obtained using HCR v3.0 is retained using HCR-Immuno. Unimodality was confirmed by Hartigan's dip test (HCR v3.0: dip = 0.0361, p = 0.602; HCR-Immuno: dip = 0.0426, p = 0.272). Puncta area is shown in pixels squared (px²).

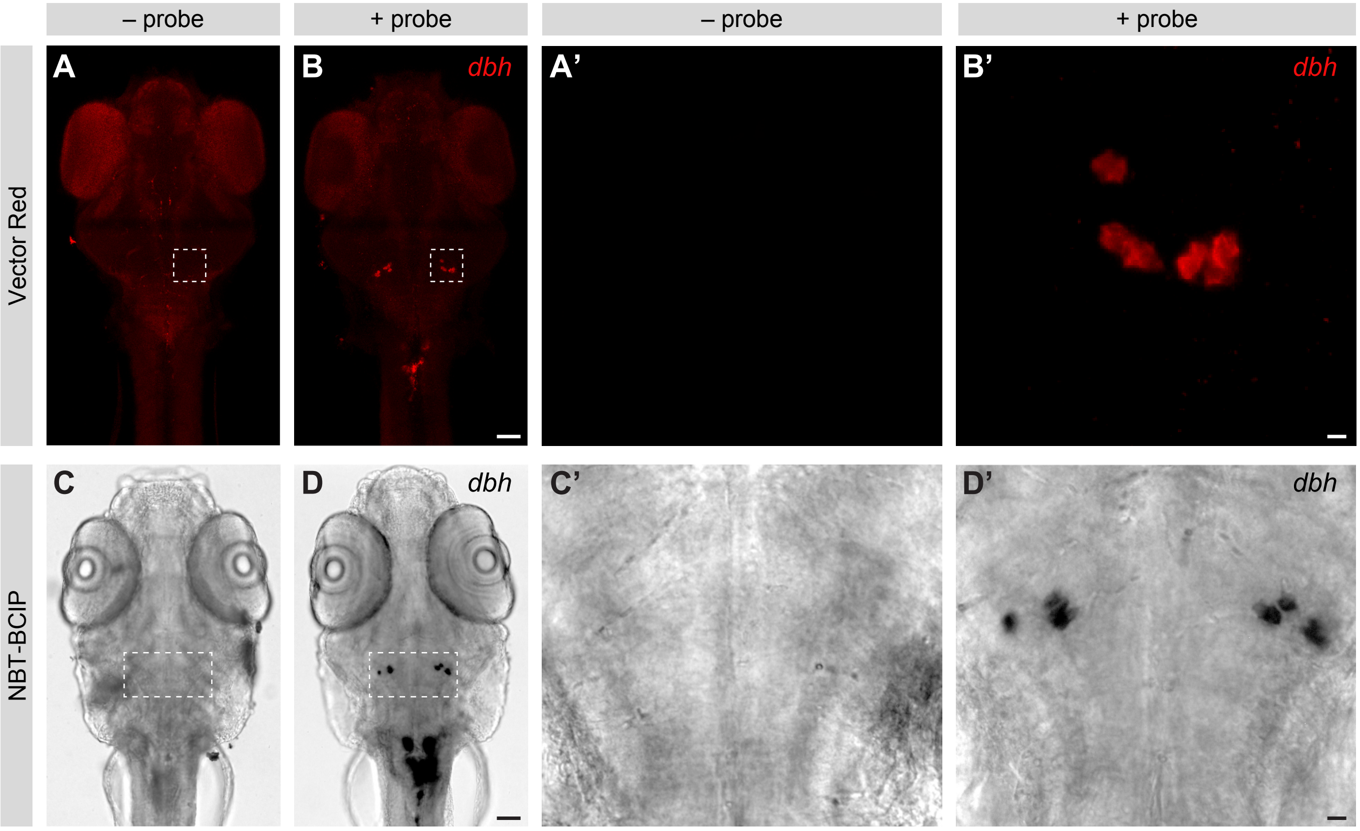


**Figure S8: HCR-Cat is compatible with alkaline phosphatase-mediated chromogenic detection.** **(A–B)** HCR-Cat using Vector Red as the alkaline phosphatase (AP) substrate in the absence of probes (**A,A'**) or following hybridization with the 20-probe *dbh* probe set (**B,B'**). Boxed regions in (**A**,**B**) are shown at higher magnification in (**A′,B′**). **(C–D)** HCR-Cat using NBT/BCIP as the AP substrate in the absence of probes (**C,C'**) or following hybridization with the 20-probe *dbh* probe set (**D, D'**). Boxed regions in (**C**,**D**) are shown at higher magnification in (**C′**,**D′**). Robust labeling of *dbh* neurons was observed with both AP substrates in probe-containing samples, whereas no signal was detected in the absence of probes. Representative images shown are maximum intensity projections of z-stacks across the entire *dbh* locus coeruleus cell populations. Vector Red fluorescence (**A–B'**) and NBT/BCIP brightfield (**C–D'**) were used. Scale bars: 100 µm **(B, D)**, 10 µm **(B′)**, and 20 µm **(D′)**.

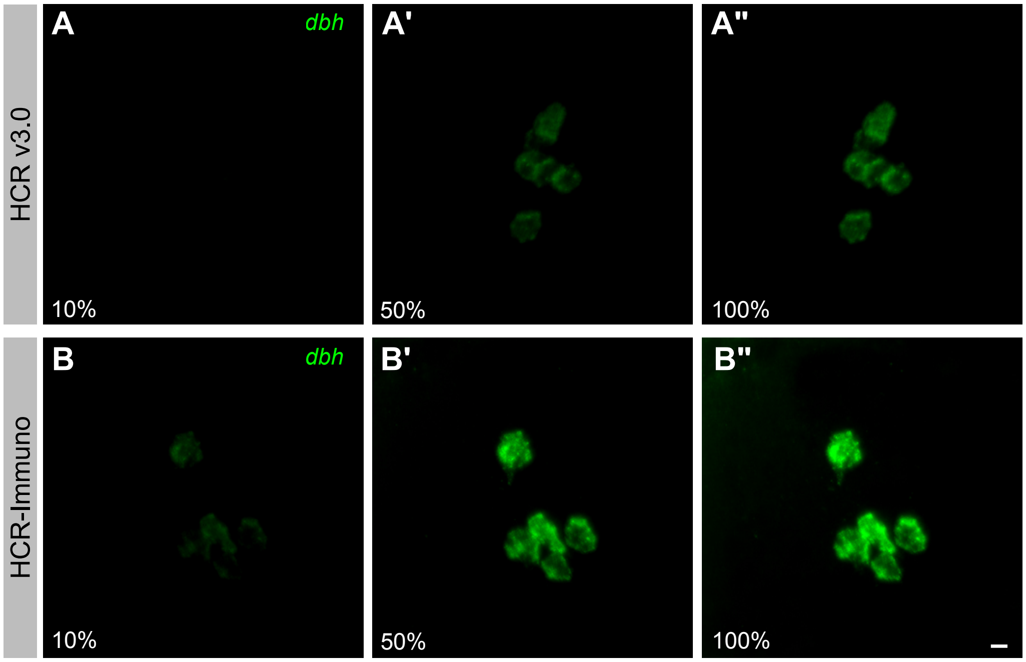


**Figure S9: HCR-Immuno is compatible with commercially available Alexa Fluor-conjugated HCR amplifier hairpins.** Representative confocal images of *dbh* neurons in the larval zebrafish locus coeruleus detected using HCR v3.0 (**A-A”**) or HCR-Immuno performed with commercially available Alexa Fluor 488-conjugated HCR amplifier hairpins and an anti-Alexa Fluor 488 antibody (**B-B”**). Images were acquired using identical imaging settings at 10%, 50%, and 100% laser power. HCR-Immuno produced substantially greater fluorescence signal than HCR v3.0 while using the same fluorophore-conjugated HCR amplifier hairpins, demonstrating that the method is compatible with existing HCR reagents without requiring custom hapten-conjugated hairpins. Scale bar: 10 μm.

**Supplementary Tables**

**Table S1. Statistical tests and results.**

**Table S2. Mouse HCR probe sequences.**
